# An *in vivo* inflammatory loop potentiates KRAS blockade

**DOI:** 10.1101/629139

**Authors:** Kristina A.M. Arendt, Giannoula Ntaliarda, Vasileios Armenis, Danai Kati, Christin Henning, Georgia A. Giotopoulou, Mario A.A. Pepe, Laura V. Klotz, Anne-Sophie Lamort, Rudolf A. Hatz, Sebastian Kobold, Georgios T. Stathopoulos

## Abstract

*KRAS* inhibitors perform inferior to other targeted drugs. To investigate a possible reason for this, we treated cancer cells with KRAS inhibitors deltarasin (targeting phosphodiesterase-δ), cysmethynil (targeting isoprenylcysteine carboxylmethyltransferase), and AA12 (targeting KRAS^G12C^), and silenced/overexpressed mutant KRAS using custom vectors. We show that *KRAS*-mutant tumor cells exclusively respond to KRAS blockade *in vivo*, because the oncogene co-opts host myeloid cells via a C-C-motif chemokine ligand 2/interleukin-1β signaling loop for sustained tumorigenicity. Indeed, *KRAS*-mutant tumors did not respond to deltarasin in *Ccr2* and *Il1b* gene-deficient mice, but were deltarasin-sensitive in wild-type and *Ccr2*-deficient mice adoptively transplanted with wild-type murine bone marrow. A KRAS-dependent pro-inflammatory transcriptome was prominent in human cancers with high *KRAS* mutation prevalence and predicted poor survival. Hence the findings support that *in vitro* systems are suboptimal for anti-KRAS drug screens, and suggest that interleukin-1β blockade might be specific for *KRAS*-mutant cancers.

## INTRODUCTION

Since its discovery, the KRAS proto-oncogene GTPase (encoded by the human *KRAS* and the murine *Kras* genes) has become the holy grail of anticancer therapy (***Esposito et al., 2019***; ***Downward, 2003***). The KRAS oncoprotein possesses a unique molecular structure that potentiates it as a driver of multiple cancer cell hallmarks (including proliferation, migration, metastasis, angiogenesis, inflammation, and apoptosis evasion), but also renders it non-actionable due to the absence of a druggable deep pocket (***Downward, 2003***;***Stephen et al., 2014***). KRAS point mutations that constitutively activate GTPase function occur most frequently in codons 12, 13,and 61 and are particularly frequent in pancreatic (70%), colorectal (35%), and lung (20%) adenocarcinomas (***Stephen et al., 2014***;***Tate et al., 2019***). However, full KRAS GTPase activity and downstream signaling additionally prerequisites its integration into the cell membrane, which is facilitated by post-translational lipidation and membrane transport of KRAS by various enzymes such as farnesyltransferase (FT), geranylgeranytransferase (GGT), isoprenylcysteine carboxylmethyltransferase (ICMT), phosphodiesteraseδ), and others (***Stephen et al., 2014***;***Simanshu et al., 2017***). To this end, therapeutic attempts to inhibit KRAS lipidation by targeting FT/GGT/ICMT were recently coupled by the development of PDEδ blockers and of allosteric and covalent inhibitors of mutated KRAS^G12C^ (***Winter-Vann et al., 2005***; ***Zimmermann et al., 2013***; ***Ostrem et al., 2013***).

Despite coordinated efforts (***Esposito et al., 2019***), anti-KRAS drug discovery is lagging behind other oncogene targets (***Stephen et al., 2014***). In addition to molecular structural considerations (***Simanshu et al., 2017***), the mode of KRAS oncogenic functions could be a reason for this. To this end, Janes and collaborators recently reported a discordance between the *in vitro* and the *in vivo* effects of a newly developed covalent KRAS^G12C^ inhibitor (***Janeset al., 2018***). This observation is relevant to other reports describing how KRAS-dependence is linked to signatures of intravital-restricted processes like inflammation and epithelial-to-mesenchymal transition (***Singh et al., 2009***; ***Sparmann and Bar-Sagi, 2004***; ***McDonald et al., 2017***) and how pro-inflammatory properties of *KRAS* mutations potentiate malignant pleural effusions in mice (***Agalioti et al., 2017***; ***Marazioti et al., 2018***).

Here we hypothesized that KRAS effects and druggability are preferentially at play *in vivo*. We tested the efficacy of three different KRAS inhibitors with divergent modes of action *in vitro* and *in vivo* using a battery of 30 natural and transduced human and murine cancer cell lines and four different methods to integrally assess tumor cell growth. We consistently show that KRAS inhibitors exert ubiquitous *in vitro* effects irrespective of cellular *KRAS* mutation status, but are specifically effective against *KRAS*-mutant tumors *in vivo*. Using transcriptome analyses of cell lines expressing endogenous or exogenous wild-type or mutant *Kras* alleles,*Ccr2* and *Il1b* gene-deficient mice, as well as adoptive bone marrow transfer, we show that mutant *Kras* establishes a proinflammatory C-C motif chemokine ligand 2 (CCL2)/interleukin-1β (IL-1β signaling loop to host myeloid cells *in vivo*, which is required for *KRAS*-mediated tumorigenicity, but also for specific KRAS inhibitor efficacy. The *KRAS/CCL2/IL1B* transcript signature is further shown to be enriched in human tumors with high *KRAS* mutation frequencies and to portend poor survival. Our data show that intact inflammatory tumor-to-host interactions are required for full KRAS inhibitor efficacy and imply that *in vitro* drug screens might not be optimal for KRAS inhibitor discovery.

## RESULTS

### Mutation-independent effects of KRAS inhibitors *in vitro*

We initially investigated the cellular responses of a battery of human and murine cell lines with known *KRAS/Kras* mutation status (***Giopanou et al., 2016***;***Agalioti et al., 2017***; ***Giannou et al., 2017***; ***Marazioti et al., 2018***;***Tate et al., 2019***; ***Kanellakis et al., 2019***) to three preclinical KRAS inhibitors: deltarasin targeting PDEδ (***Zimmermann et al., 2013***), AA12 allosterically targeting KRAS^G12C^(***Ostrem et al., 2013***), and cysmethynil targeting ICMT (***Winter-Vann et al., 2005***) (***Figure 1A*** and ***Figure 1–figure supplement 1***). For this, widely used assays were employed based on literature searches (***Figure 1–figure supplement 2*** and ***Figure 1–source data 1***).

**Figure 1.**
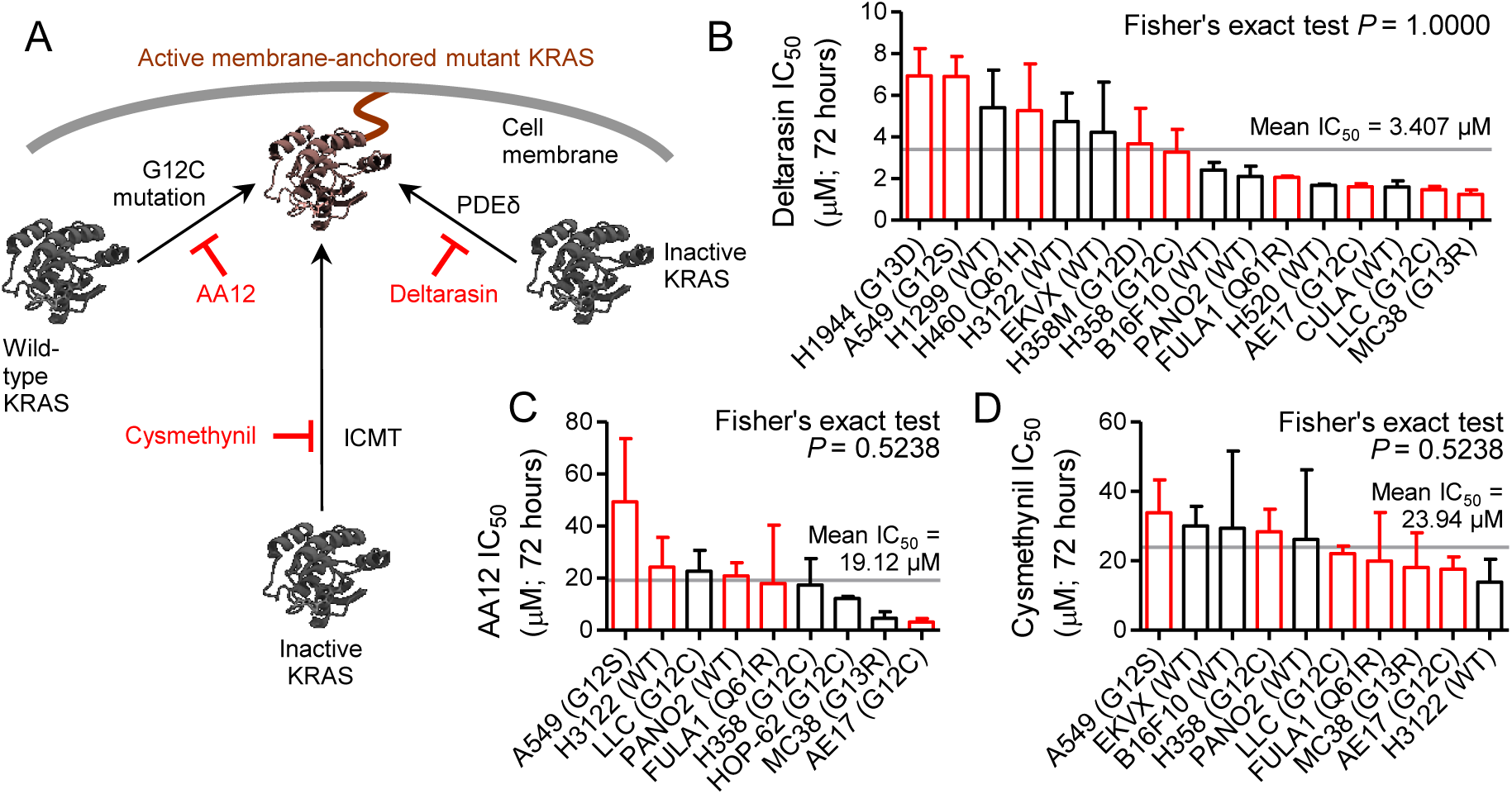
Pharmacologic evidence for *KRAS* mutation-independence *in vitro*. Different mouse and human tumor cell lines with (red) and without (black) *Kras/KRAS* mutations (codon changes are given in parentheses) were assessed for cell viability by colorimetric WST-8-assay after 72-hour treatments with three different KRAS inhibitors (*n* = 3/data-point). **(A)** Graphical abstract showing molecular targets of preclinical KRAS inhibitors AA12, cysmethynil, and deltarasin. **(B-D)** Fifty percent inhibitory concentrations (IC_50_) of deltarasin (B), AA12 (C), and cysmethynil (D) by WST-8 assay. Data presented as mean ± SD. Grey lines represent the mean of all cell lines tested, which was used to dichotomize cell lines into sensitive and resistant. *P*, probability by Fisher’s exact test for cross-tabulation of *Kras/KRAS* mutation status to drug sensitivity/resistance. KRAS, KRAS proto-oncogene GTPase; WT, wild-type.

Initially, 50% inhibitory concentration (IC_50_) values were calculated from water soluble tetrazolium-8 (WST-8) assays doneafter 72 hours of treatment with half-log-incremental drug concentrations. Unexpectedly, all three KRAS inhibitors showed comparable modest *in vitro* efficacy across all cell lines tested, independent of their *KRAS/Kras* mutation status, with IC_50_ values between 1-50 µM (***Figures 1B-D***,***Tables 1-3***, ***Figure 1– figure supplement 3***, and ***Figure 1–source data2***). A literature search revealed that this was generally true for developmental KRAS inhibitors compared with tyrosine kinase inhibitors (***Figure 1– figure supplement 4*** and ***Figure 1–source data3***). To extend these results, we analyzed the response of eight selected murine and human cell lines to 60% inhibitory concentrations (IC_60_) of deltarasin in an *in vitro* colony formation assay. Again, deltarasin efficacy was independent of *KRAS/Kras* mutation status (***Figures 2A and B***, ***Figure 2–figure supplement 1***, and ***Figure 2–source data 1***). Since KRAS activates the mitogen-activated protein kinase cascade inducing phosphorylation of extracellular-signal regulated kinase (ERK), we quantified total (t)- and phospho (p)-ERK relative to glyceraldehyde 3-phosphate dehydrogenase (GAPDH) in 12 murine and human cell lines treated with saline or IC_60_deltarasin. In line with the above, deltarasin inhibited p-ERK independent from cellular *KRAS/Kras* mutation status (***Figure 2C and D***, ***Figure 2–figure supplement 2***, and ***Figure 2–source data 2***). Thus, pharmacologic KRAS inhibition does not reveal KRAS-dependence *in vitro*.

**Figure 2.**
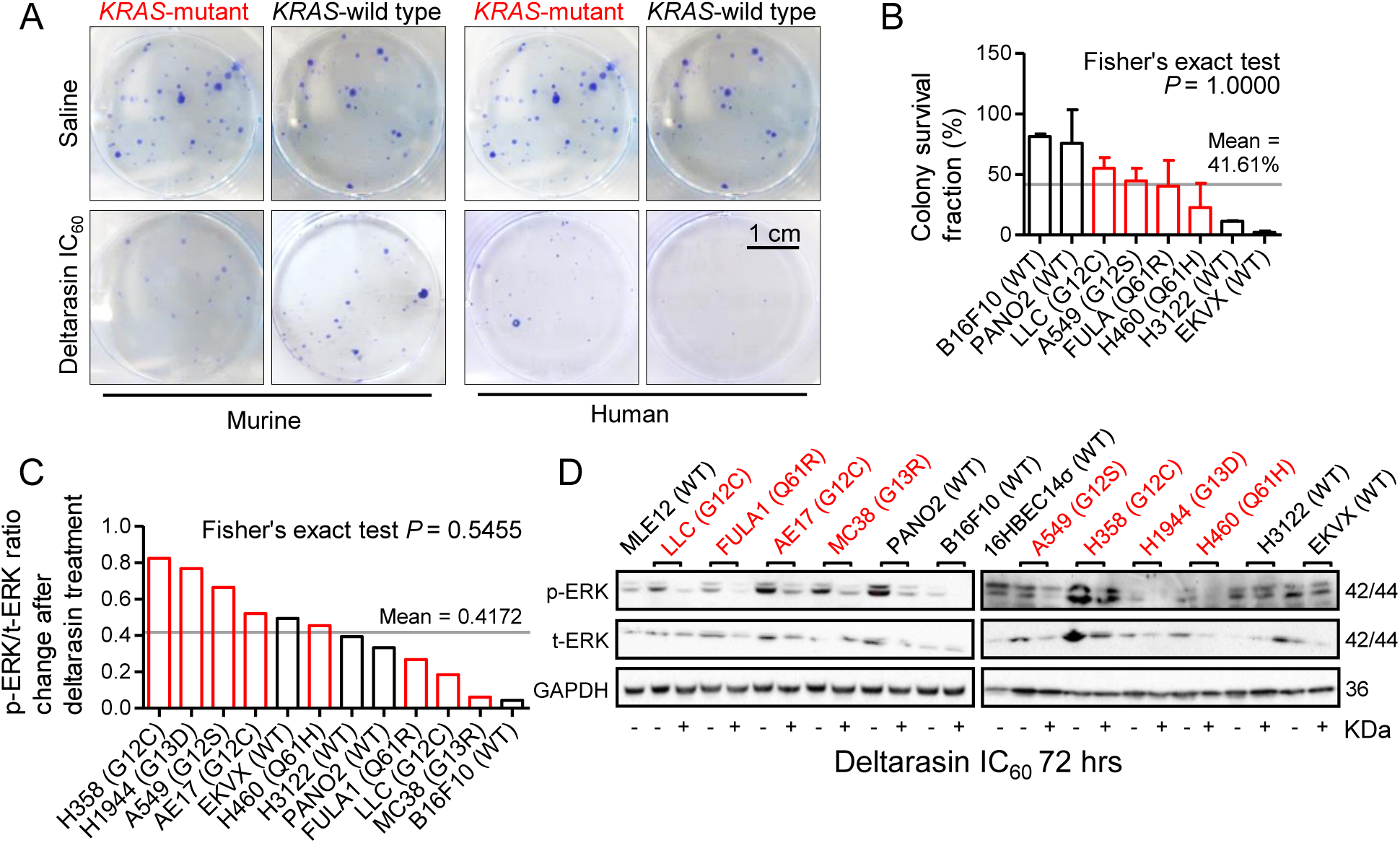
*KRAS* mutation-independence ofcolony formation and ERK phosphorylation. Different mouse and human tumor cell lines with (red) and without (black) *Kras/KRAS* mutations (codon changes are given in parentheses) were assessed for colony formation by crystal violet-stained colony counts and for ERK phosphorylation by phospho (p)- and total (t)- ERK immunoblots after 72-hour treatments with the KRAS inhibitor deltarasin (*n* = 3/data-point). **(A, B)** Representative images of colonies after saline or IC_60_ deltarasin treatment (A) and colony survival fraction (B) after IC_60_ deltarasin normalized to saline treatment.**(C, D)** Quantification of normalized p-ERK/t-ERK signal change after IC_60_deltarasin normalized to saline treatment (C) and representative immunoblots (D). **(B, C)** Data presented as mean ± SD. Grey lines represent the mean of all cell lines tested, which was used to dichotomize cell lines into sensitive and resistant. *P*, probability by Fisher’s exact test for cross-tabulation of *Kras/KRAS* mutation status to drug sensitivity/resistance. KRAS, KRAS proto-oncogene GTPase; WT, wild-type; GAPDH, glyceraldehyde 3-phosphate dehydrogenase.

### Specific *in vivo* effects of deltarasin against *KRAS*-mutant tumors

To replicate these results *in vivo*, we induced subcutaneous (sc) tumors in *C57BL/6*, *FVB*,and *Rag2^-/-^*mice, as appropriate, using six different cancer cell lines. After tumor establishment, which was diagnosed when both tumor volume superseded 100 mm^3^ and tumor latency 14 days post-sc injection, we initiated daily intraperitoneal (ip) saline 2% dimethyl sulfoxide (DMSO) (hereafter referred to as saline) or deltarasin (15 mg/Kg in saline 2% DMSO, hereafter referred to as deltarasin) treatments. Interestingly, deltarasin selectively inhibited the sc growth of murine and human *KRAS*-mutant (*KRAS*^MUT^) tumors, but had no effect on *KRAS*-wild-type (*KRAS*^WT^) tumors (***Figures 3A and B*** and ***Figure 3–source data 1***). Moreover, forced overexpression of a custom-made plasmid encoding *Kras*^G12C^ (p*Kras*^G12C^) in *KRAS*^WT^ mouse and human cancer cells accelerated tumor growth and restored the response to the drug over forced overexpression of empty vector (p*C*) (***Figure 3C***,***Figure 3–figure supplement 1***, and ***Figure 3–source data 2***). Taken together, these data show that deltarasin-mediated KRAS inhibition selectively halts the growth of *KRAS*^MUT^ cancer cells *in vivo*.

**Figure 3.**
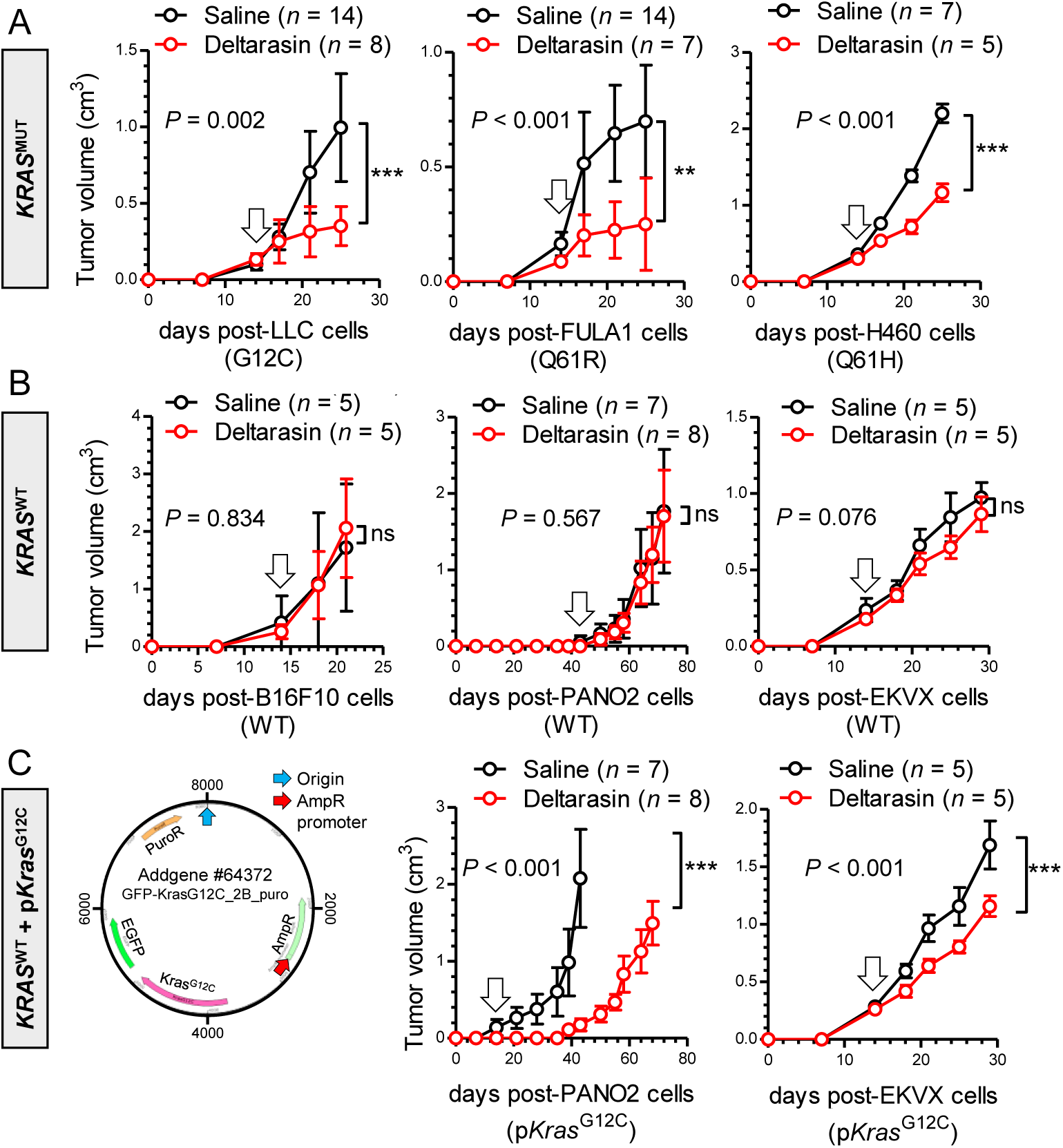
Deltarasin-mediated demonstration of *KRAS* mutation-dependence *in vivo*. Different mouse and human tumor cell lines with (**A**; *KRAS*^MUT^) and without (**B**; *KRAS*^WT^) endogenous *Kras/KRAS* mutations (codon changes are given in parentheses), as well as *KRAS*^WT^ cell lines forcedly expressing a plasmid encoding mutant murine *Kras*^G12C^ (**C**; p*Kras*^G12C^), were injected into the rear flank (10^6^ tumor cells sc) of *C57BL/6* (LLC, B16F10, and PANO2 cells), *FVB* (FULA1 cells), or *Rag2^-/-^* (H460 and EKVX cells) mice. After tumor establishment (tumor volume > 100 mm^3^ and latency > 14 days; arrows), mice were randomly allocated to daily ip L saline 2% DMSO (black) or 15 mg/ Kg deltarasin in 100 μ DMSO (red). Tumor growth was assessed by measuring three vertical tumor dimensions. Data presented as mean ± SD.*n*, sample size; *P*, overall probability, 2-way ANOVA; ns,**, and ***: *P* > 0.05, *P* < 0.01, and *P* <0.001, respectively, Bonferroni post-test.

### Genetic*KRAS*manipulation reveals *in vivo-*restricted *KRAS*-dependence

To further validate the observed *in vivo*-restricted specificity of deltarasin, we overexpressed random (sh*C*) or anti-*Kras*-specific shRNA (sh*Kras*) in *Kras*^MUT^ parental cell lines or p*Kras*^G12C^in *Kras*^WT^ parental cell lines (***Agalioti et al., 2017***). In accord with pharmacologic

KRAS inhibition, genetic *Kras* modulation did not impact the *in vitro* response of cancer cell lines to deltarasin, as determined by WST-8IC_50_ values and ERK activation levels (***Figure 4***,***Figure 4–figure supplement 1***, and ***Figure 4–source data 1***). In contrast to the lack of *Kras*-dependence *in vitro*, mutant *Kras* was required and sufficient for sustained tumor growth *in vivo* (***Figure 5*** and ***Figure 5–source data 1***): murine cell lines expressing sh*Kras* displayed statistically (*P* < 0.001) and biologically (50-90% inhibition) significantly decreased tumor growth compared with parental cell lines expressing sh*C*. *Vice versa*, p*Kras*^G12C^ overexpression accelerated tumor growth compared with overexpression of p*C*. Collectively, these results support that, similar to drug-based KRAS inhibition, genetic *Kras* modulation selectively impacts tumor growth *in vivo*.

**Figure 4.**
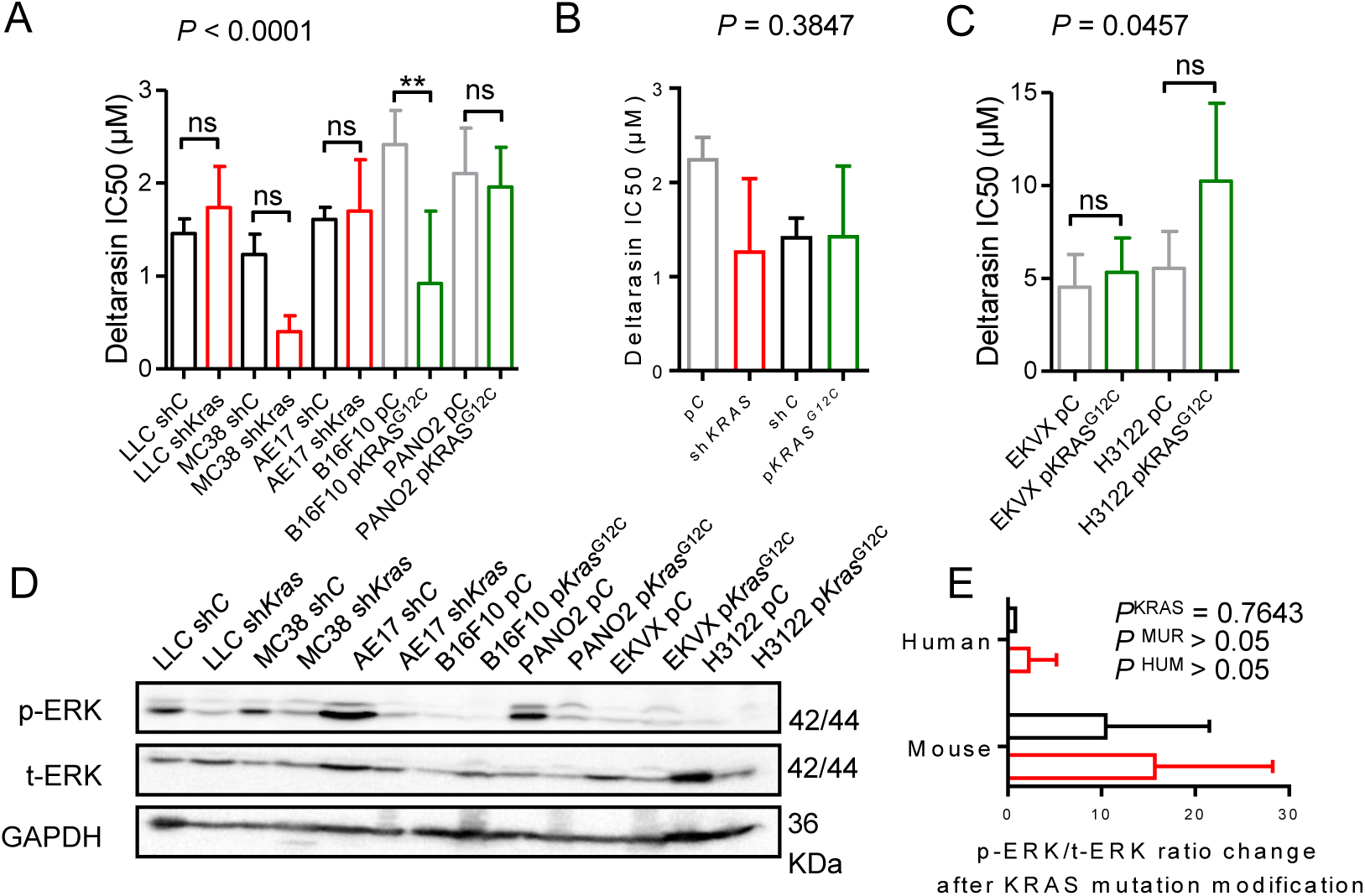
Genetic evidence for *KRAS* mutation-independence *in vitro*. (A) Different murine parental (black/grey: stably expressing random shRNA, sh*C*, or control plasmid, p*C*) or *Kras*-modified (red: stably expressing sh*Kras*; green: stably expressing mutant *Kras*^G12C^ plasmid, p*Kras*^G12C^) tumor cell lines were assessed for cell viability (IC_50_byWST-8-assay; *n*= 3/data-point) after 72 hours of deltarasin treatment. **(B)** Summary of averaged deltarasin IC_50_ values from all cell lines from (A) (*n* = 3 cell lines/group). **(C)** Human parental (black/grey: stably expressing control plasmid p*C*) or *KRAS*-modified (green: stably expressing p*Kras*^G12C^) tumor cell lines were assessed for cell viability by WST-8assay (*n* = 3/data-point) after 72 hours of deltarasin treatment. **(D)** Immunoblots of cell lines from (A) for p-ERK, t-ERK and GAPDH. **(E)** Quantification of normalized p-ERK/t-ERK signal from (D). Data were summarized by mutation status and origin. *P*, overall probability by one-way (A-C) and two-way (E) ANOVA. ns and **: *P* > 0.05 and *P* < 0.01, respectively, for the indicated comparisons by Bonferroni post-tests. Data are presented as mean ± SD.

**Figure 5.**
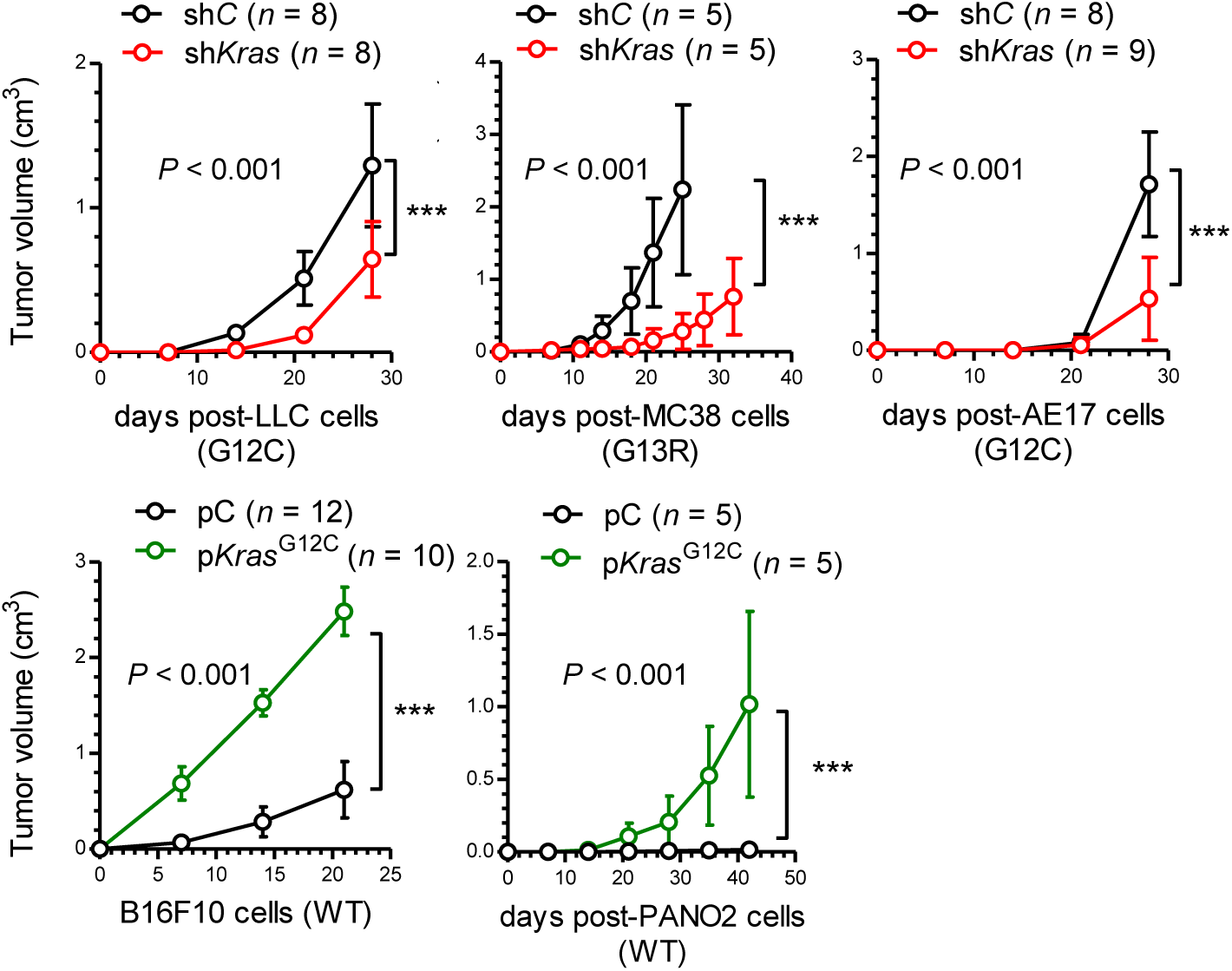
Genetic manipulation of *Kras* reveals *in vivo*-restricted KRAS dependence. Different murine parental (black/grey: stably expressing random shRNA, sh*C*, or control plasmid, p*C*) or *Kras*-modified (red: stably expressing sh*Kras*; green: stably expressing mutant *Kras*^G12C^ plasmid, p*Kras*^G12C^) tumor cell lines were injected into the rear flank (10^6^ tumor cells sc) of *C57BL/6* mice for induction of flank tumors by genetically modified cells (red, sh*Kras*; green, p*Kras*^G12C^) or control cells (black, sh*C* or p*C*). *P*, overall probability by two-way ANOVA. ***: *P*< 0.001 for the indicated comparisons by Bonferroni post-tests. Data are presented as mean ± SD.

### A mutant *Kras* transcriptome signature contains *Ccl2* and *Il1r1*

In an effort to identify *Kras*^MUT^-driven genes responsible for *in vivo* restricted KRAS-dependence, we analyzed the global transcriptomes of the parental and *Kras*-modulated murine cell lines described above and of benign samples [whole lungs, tracheal epithelial cells (TEC), and bone marrow-derived macrophages (BMDM) and mast cells (BMMC); GEO dataset GSE130624]. Unsupervised hierarchical clustering showed an absolute segregation of benign, *Kras*^WT^, and *Kras*^MUT^ samples by 1408 differentially expressed genes (ΔGE) using an ANOVA *P* < 0.05 threshold (***Figure 6A***). Paired analyses of five isogenic cancer cell line doublets with modulated *Kras* (LLC, MC38, and AE17 cells expressing sh*C* versus sh*Kras* and PANO2 and B16F10 cells expressing p*C* versus p*Kras*^G12C^) identified another 3432 *Kras*-responsive transcripts. Out of the 170 transcripts that were present in both gene sets, 42 were both differentially represented in benign, *Kras*^WT^, and *Kras*^MUT^ samples and responsive (ΔGE > 1.40) to *Kras* modulation, including *Kras per se* (***Figure 6B*** and ***Figure 6–source data 1***).

**Figure 6.**
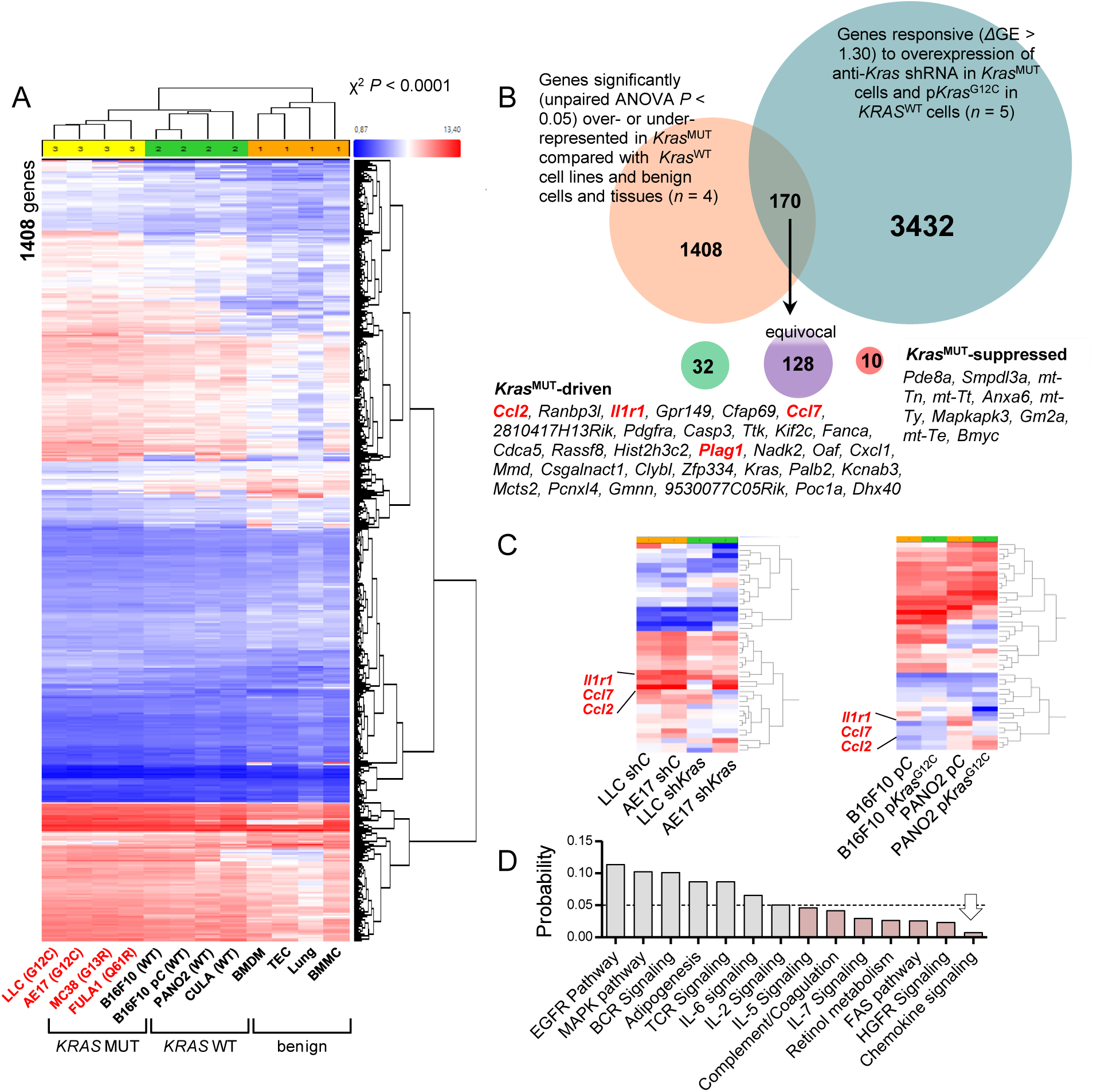
A 42-gene inflammatory signature of KRAS-dependence. (A) **Unsupervised** hierarchical clustering of gene expression of *Kras*-mutantand *Kras*-*WT* cancer cell lines, as well as benign cells and tissues. **(B)** Venn diagram of analytical strategy of transcriptome analysis. **(C)** Unsupervised hierarchical clustering of gene expression of *Kras*-modified cancer cell line doublets reveals co-clustering of *Il1r1* and *Ccl2*. **(D)** WikiPathway analysis showing pathways significantly overrepresented in the *KRAS* signature.

Interestingly, *Il1r1*, *Ccl7*, and *Ccl2* were among those genes and clustered tightly together (***Figure 6C***) and chemokine signaling was the pathway most significantly perturbed by *Kras* modulation on WikiPathway analysis (***Kelder et al., 2012***) (***Figure 6D*** and ***Figure 6–source data 2***). We next translated our 42-gene murine mutant *Kras* signature to their 37 human orthologues using OrthoDB (https://www.orthodb.org/; ***Kriventseva et al., 2019***) and ran gene set enrichment analyses (GSEA;http://software.broadinstitute.org/gsea/index.jsp; ***Subramanian et al., 2005***). Interestingly, our humanized *KRAS*^MUT^ signature was enriched in only two out of the Broad Institute’s 50 hallmark signatures: positively in the signature “inflammatory response” and negatively in the signature “G2M-checkpoint” (***Figure 7A***). Moreover, this mutant *KRAS* signature was significantly positively enriched in *KRAS*-versus *EGFR*-mutant lung adenocarcinomas (LADC) from the BATTLE trial (GEO dataset GSE31852; ***Kim et al., 2011***; ***Kabbout et al., 2013***) (***Figure 7B***). In this connection, we recently reported that mutant *KRAS* drives C-C-motif chemokine ligand 2 (CCL2) and interleukin-1 receptor 1 (Il1R1) expression to establish inflammatory feedback loops with interleukin-1β (IL-1β malignant pleural effusions and developing *KRAS*^MUT^ LADC (***Giannou et al., 2015***; ***Agalioti et al., 2017***; ***Marazioti et al., 2018***; ***Lilis et al., 2019***). Collectively, the data suggested that *in vivo*-restricted *KRAS*^MUT^-dependence might be mediated by proinflammatory signals to CCR2+ IL-1β-secreting host cells.

**Figure 7.**
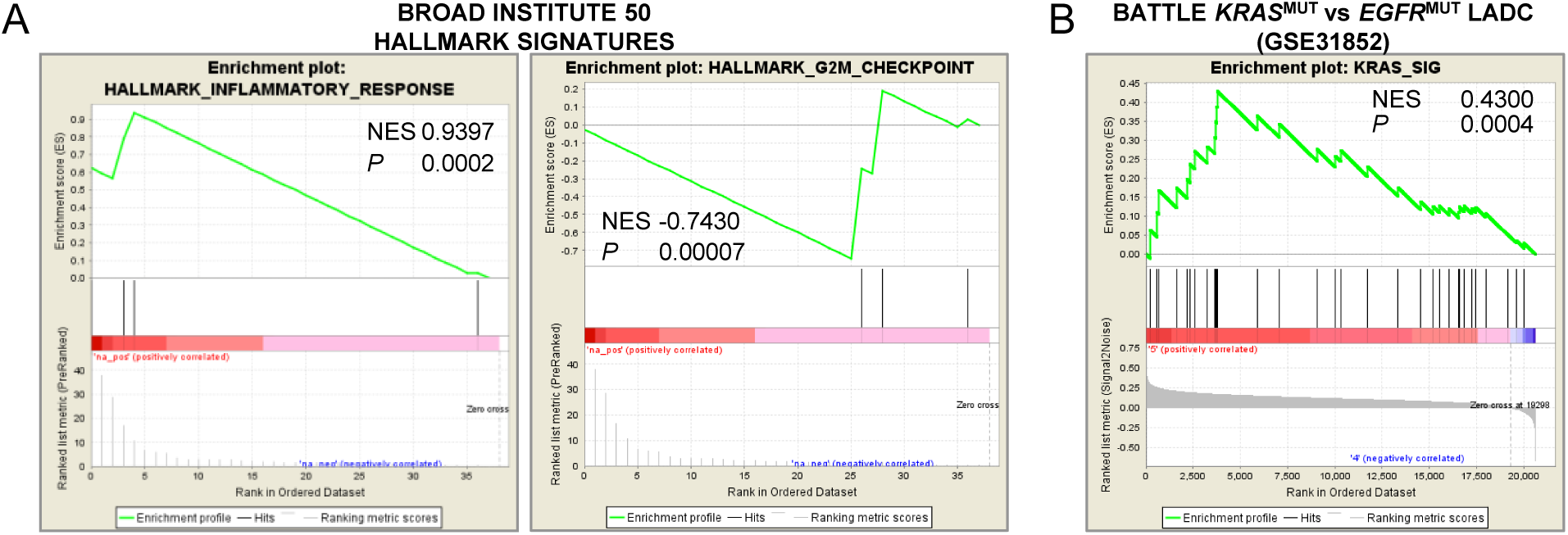
Enrichment of the murine KRAS-dependence signature in human transcriptomes. GSEA of 37 human orthologues of the murine *KRAS* signature against the Broad Institute’s 50 hallmark signatures showing positive enrichment in the “inflammatory response” and negative enrichment in the “G2M checkpoint” signatures **(A)** and against KRAS- (*n* = 21) versus EGFR- (*n* = 17) -mutant lung adenocarcinomas (LADC) from BATTLE **(B)** revealing positive enrichment of our KRAS signature in human *KRAS*-mutant LADC. NES, normalized enrichment score; *P*, family-wise error rate probability.

### CCR2+ IL-1**β**-secreting myeloid cells potentiate *in vivo* KRAS-dependence

These results led us to the hypothesis that CCR2+ IL-1β-secreting myeloid cells are required for *in vivo* KRAS-dependence (***Figure 8A***). Indeed, numerous such cells co-expressing CCR2 and IL-1β were identified in the stromata of our experimental *KRAS*-mutant tumors by immunohistochemistry (***Figure 8B***). To definitively test our hypothesis, we induced flank tumors by injecting one million LLC cells (*Kras*^G12C^) sc into syngeneic *C57BL/6*mice competent (*WT*) or deficient (*Il1b^-/-^, Ccr2^-/-^*) in the *Il1b* and *Ccr2* genes (***Boring et al., 1997***; ***Horai et al., 1998***). Mice haplo/diplo-insufficient in the *Cxcr1* and *Cxcr2* chemokine receptor genes (*Cxcr1^-/-^, Cxcr2^+/-^*) (***Cacalano et al., 1994***; ***Sakai et al., 2011***; ***Giannou et al., 2017***) were also employed as additional controls for *Ccr2^-/-^* mice and daily ip saline (2% DMSO) or 15 mg/Kg deltarasin (in saline 2% DMSO) treatments were initiated when tumors reached 100 mm^3^ volumes and 14 days latency, as above. Expectedly, deltarasin treatment statistically and biologically significantly inhibited LLC tumor growth in *WT*, *Cxcr1^-/-^*, and *Cxcr2^+/-^* mice. However, deltarasin effects were diminished in *Il1b^-/-^* and completely abrogated in *Ccr2^-/-^* mice (***Figure 8C*** and ***Figure 8– source data 1***). To exclude the possibility of developmental effects of knockout mice, we total-body irradiated (900 Rad) *Ccr2^-/-^* mice and performed adoptive bone marrow transplants (BMT) from *WT* or *Ccr2^-/-^*donors, as described and validated previously (***Giannou et al., 2015***; ***Marazioti et al., 2018***). For this experiment, *WT* and *Ccr2^-/-^* mice back-crossed > F12 to the *FVB* strain were used together with syngeneic FULA1 cells (*Kras*^Q61R^) to obtain results with another cell line harboring a different *Kras* mutation and a broad mutation spectrum relevant to human *KRAS*-mutant LADC (***Kanellakis et al., 2019***). Again, daily ip saline or deltarasin treatments (all in saline 2% DMSO) were started when tumors reached > 100 mm^3^ volumes at latency > 14 days. Expectedly, *Ccr2^-/-^* chimeras receiving *Ccr2^-/-^* BMT did not respond to deltarasin treatment, but *Ccr2^-/-^* chimeras receiving *WT* BMT displayed markedly increased tumor growth as well as a statistically and biologically significant inhibition by deltarasin treatment (***Figure 9A*** and ***Figure 9–source data 1***). Collectively, these results indicate that myeloid CCR2 and IL-1β are required for deltarasin efficacy against *Kras*-mutant tumors *in vivo*.

**Figure 8.**
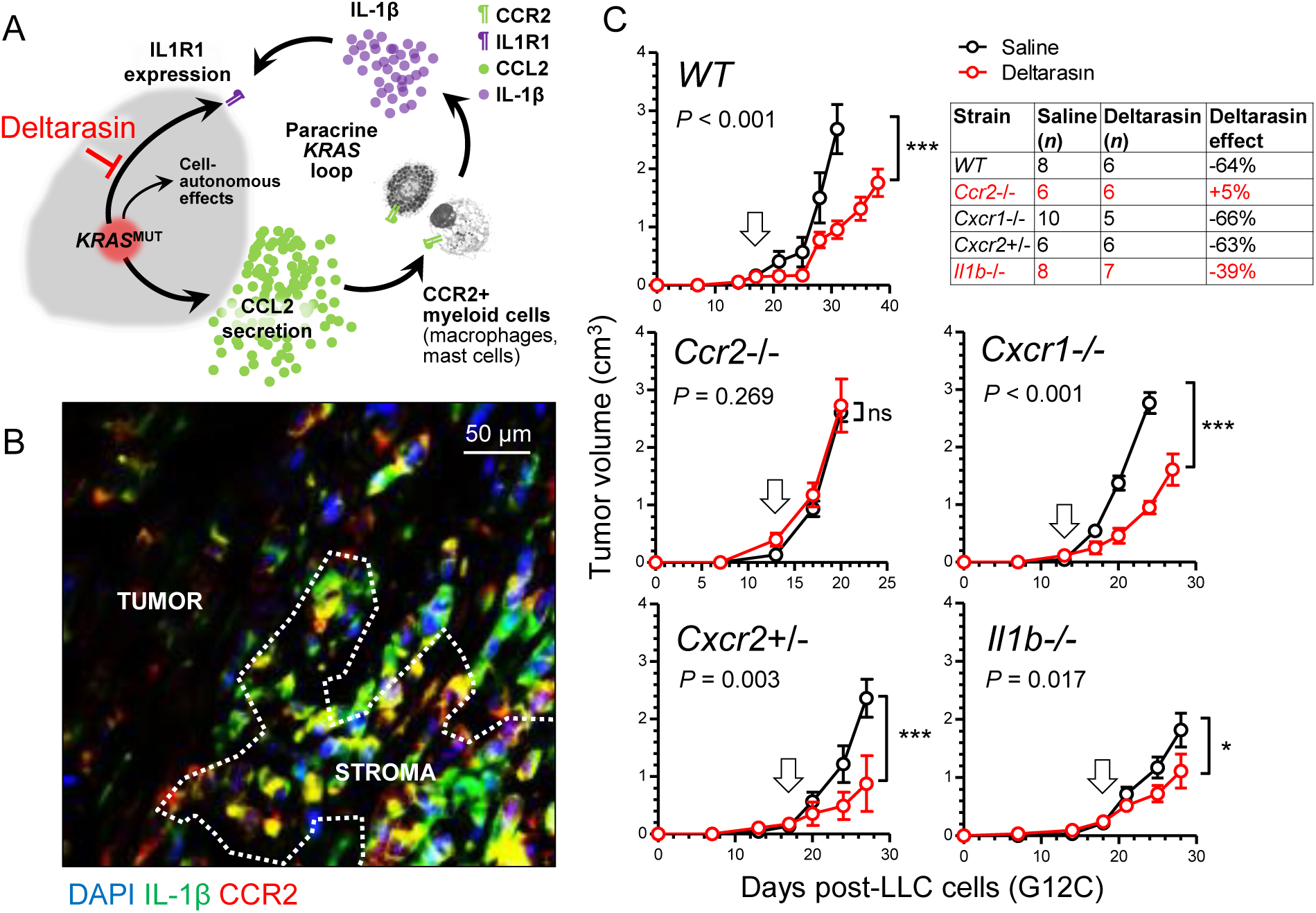
A requirement for host *Ccr2* and *IL1b* for KRAS dependence *in vivo*. (A) Graphical abstract of the proposed mechanism of *in vivo* restricted KRAS dependence. -co-staining of a *KRAS*-mutant tumor from a *Rag2^-/-^*β mouse showing co-localization of the two proteins in the tumor stroma. Image was taken using an AxioImager.M2 (Zeiss; Jena, Germany) and a 60x objective. **(C)** Syngeneic *C57BL/6* mice competent (*WT*) or deficient (*Il1b^-/-^, Ccr2^-/-^*) in the *Il1b* and *Ccr2* genes or haplo/diplo-insufficient in the *Cxcr1* and *Cxcr2* chemokine receptor genes (*Cxcr1^-/-^, Cxcr2^+/-^*) received10^6^ LLC cells (*Kras*^G12C^) sc followed by daily ip saline 2% DMSO (black) or 15 mg/Kg deltarasin in saline 2% DMSO (red) treatments initiated when tumors reached >100 mm^3^ volumes and > 14 days latency (arrows). Data are presented as mean ± SD. *P*, overall probabilities by 2-way ANOVA; ns, *, and ***: *P*> 0.05, *P* < 0.05, and *P* < 0.001 for the indicated comparisons by Bonferroni post-tests. Table shows animal numbers used and percentile tumor inhibition by deltarasin compared with saline.

**Figure 9.**
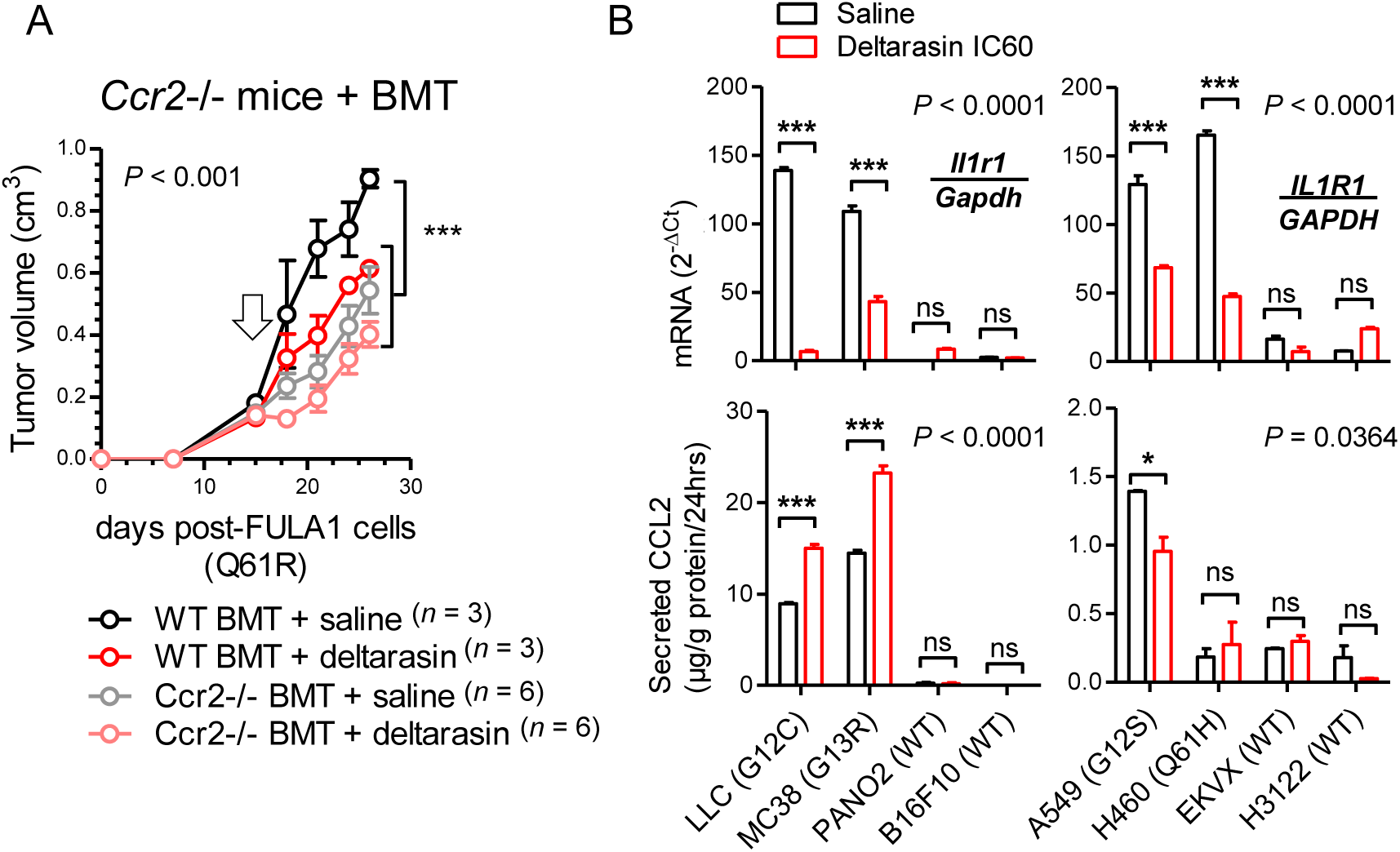
*In vivo* KRAS-dependence requires myeloid *Ccr2* and is abolished by deltarasin treatment via downregulation of *IL1R1* expression in *KRAS*-mutant cancer cells. **(A)** Total-body irradiated (900 Rad) *Ccr2^-/-^* mice received adoptive BMT from *WT* or *Ccr2^-/-^* donors (all back-crossed > F12 to the *FVB* strain). After one month allowed for chimeric bone marrow reconstitution, chimeras received 10^6^ syngeneic FULA1 cells (*Kras*^Q61R^) sc. Daily ip saline 2% DMSO or deltarasin (15 mg/Kg in saline 2% DMSO) treatments were started when tumors > 100 mm^3^ were established at > 14 days latency (arrow). Data are presented as mean ± SD. *P*, overall probabilities by 2-way ANOVA; ***: *P* < 0.001 for the indicated comparisons by Bonferroni post-tests. **(B)** *Il1r1/IL1R1* mRNA expression by qPCR (top) and CCL2 protein secretion by ELISA (bottom) of mouse (left) and human (right) cancer cell lines treated with saline 2% DMSO or deltarasin IC_60_in saline 2% DMSO for 72 hours. Data are presented as mean ± SD. *P*, overall probabilities by 2-way ANOVA; ns, *, and ***: *P* > 0.05, *P* < 0.05 and *P* < 0.001, respectively, for the indicated comparisons by Bonferroni post-tests.

### Deltarasin limits IL-1**β** sensing by *KRAS*-mutant tumor cells

We next interrogated the mechanism of *in vivo*-restricted deltarasin dependence. Based on the microarray-derived mutant *Kras* signature that encompassed *Ccl2* and *Il1r1* (***Figure 6***) and our previous reports of mutant *KRAS*-mediated transcriptional regulation of *CCL2* and *IL1R1* (***Agalioti et al., 2017***; ***Marazioti et al., 2018***), we tested whether deltarasin blocks expression of these two genes (***Figure 9B*** and ***Figure 9–source data 2***). Indeed, *Kras/KRAS*^MUT^ mouse and human cancer cell lines displayed markedly increased baseline *Il1r1/IL1R1* mRNA expression compared with *Kras/KRAS*^WT^ cell lines, and significantly downregulated *Il1r1/IL1R1* transcript levels after deltarasin treatment. On the contrary, only some *Kras/KRAS*^MUT^ cell lines displayed increased baseline CCL2 protein secretion compared with *Kras/KRAS*^WT^ cell lines, and CCL2 elaboration was not consistently blocked by deltarasin treatment, suggesting that deltarasin-mediated downregulation of *Il1r1/IL1R1* expression delivers the bulk of the drug’s *in vivo* effects.

### An inflammatory *CCL2/IL1B* signature in *KRAS*-mutant human cancers

To investigate the relevance of our findings to *KRAS*-mutant human cancers, we analyzed the average expression of *KRAS*, *CCL2*, and *IL1B* genes in another publicly available dataset from the BATTLE study (GEO dataset GSE43458; ***Kim et al., 2011***; ***Kabbout et al., 2013***). Interestingly, mean *KRAS/CCL2/IL1B* expression was statistically significantly increased in smokers’ LADC (*n* = 40) compared with never-smokers’ LADC (*n* = 40) and normal lung tissue samples (*n* = 30) (***Figure 10A*** and ***Figure 10–source data 1***). Since *KRAS* mutations are more frequent in LADC of smokers (***Campbell et al., 2016***), this finding suggested that our inflammatory signature was overrepresented in tumors with higher *KRAS* mutation frequencies. This was also true in another dataset from patients with breast, colorectal, and lung cancer (GEO dataset GSE103512; ***Brouwer-Visser et al., 2018***), where mean *KRAS/CCL2/IL1B* expression was significantly higher in lung and colorectal cancer, which have higher *KRAS* mutation rates (***Tate et al., 2019***), compared with breast cancer (***Figure 10B*** and ***Figure 10–source data 2***). Finally, online Kaplan-Meier analyses (http://www.kmplot.com; ***Győrffy et al., 2013***) using lung cancer patient data were done (***Figure 11***). These revealed that in patients with LADC (a tumor with high *KRAS* mutation frequency) high *KRAS/CCL2/IL1B* expression levels portended 93% increased odds of death regardless of smoking status. On the contrary, *KRAS/CCL2/IL1B* expression did not impact the survival of patients with squamous cell lung carcinoma (a tumor with low *KRAS* mutation frequency). When exclusively smokers were examined (thereby enriching the sample for *KRAS*-mutant patients), high *KRAS/CCL2/IL1B* expression levels portended 128% increased odds of death in LADC and continued to have no impact on the survival of patients with squamous cell lung carcinoma. Taken together, these data suggest that *KRAS/CCL2/IL1B* transcripts are overexpressed in human *KRAS*-mutant cancers and detrimentally affect survival. Moreover, the results supported that the proposed KRAS-driven inflammatory loop may be clinically relevant.

**Figure 10.**
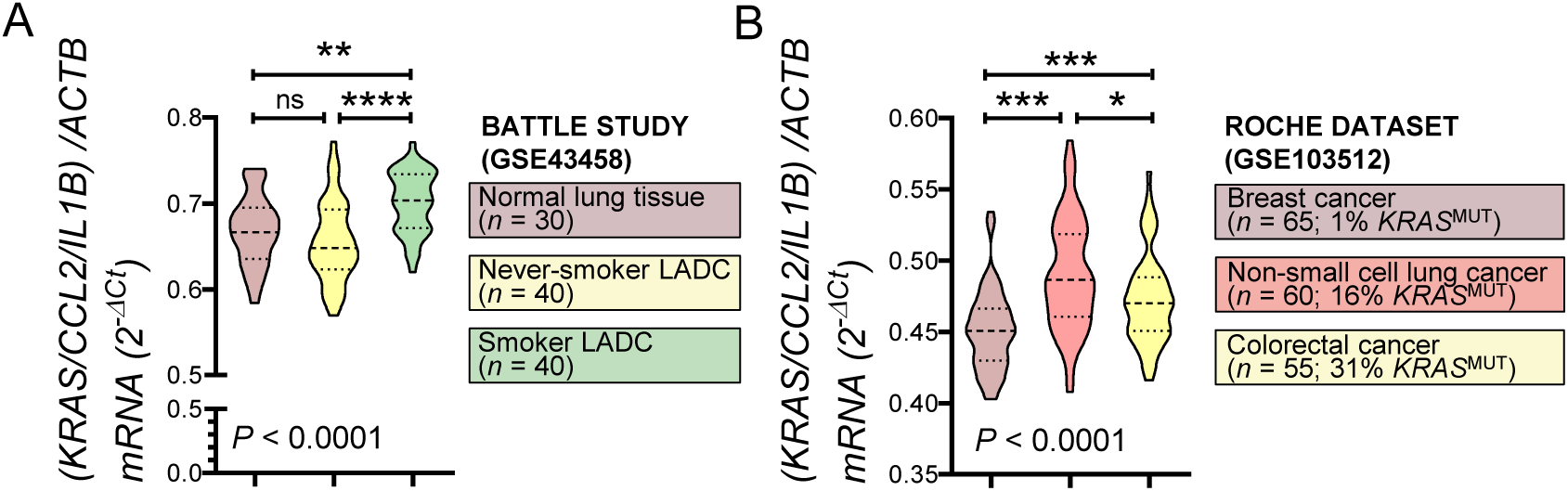
Mean expression of *KRAS/CCL2/IL1B* is increased in *KRAS*-mutant cancers. (A) Average *KRAS/CCL2/IL1B* expression normalized to *ACTB* in lung adenocarcinomas (LADC) from smokers and never-smokers and normal lung tissue from never-smokers from the BATTLE study (GSE43458). **(B)** *KRAS/CCL2/IL1B* expression normalized to *ACTB* in breast, non-small cell lung, and colorectal cancer (ROCHE study GSE103512). *KRAS* mutation frequencies of these tumor types are from COSMIC (***Tate et al., 2019***). Data are presented as violin plots. *P*, overall probability by one-way ANOVA. ns, *, **, and ***: *P* > 0.05, *P* < 0.05,*P* < 0.01, and *P* < 0.001, respectively, for the indicated comparisons by Bonferroni post-tests.

**Figure 11.**
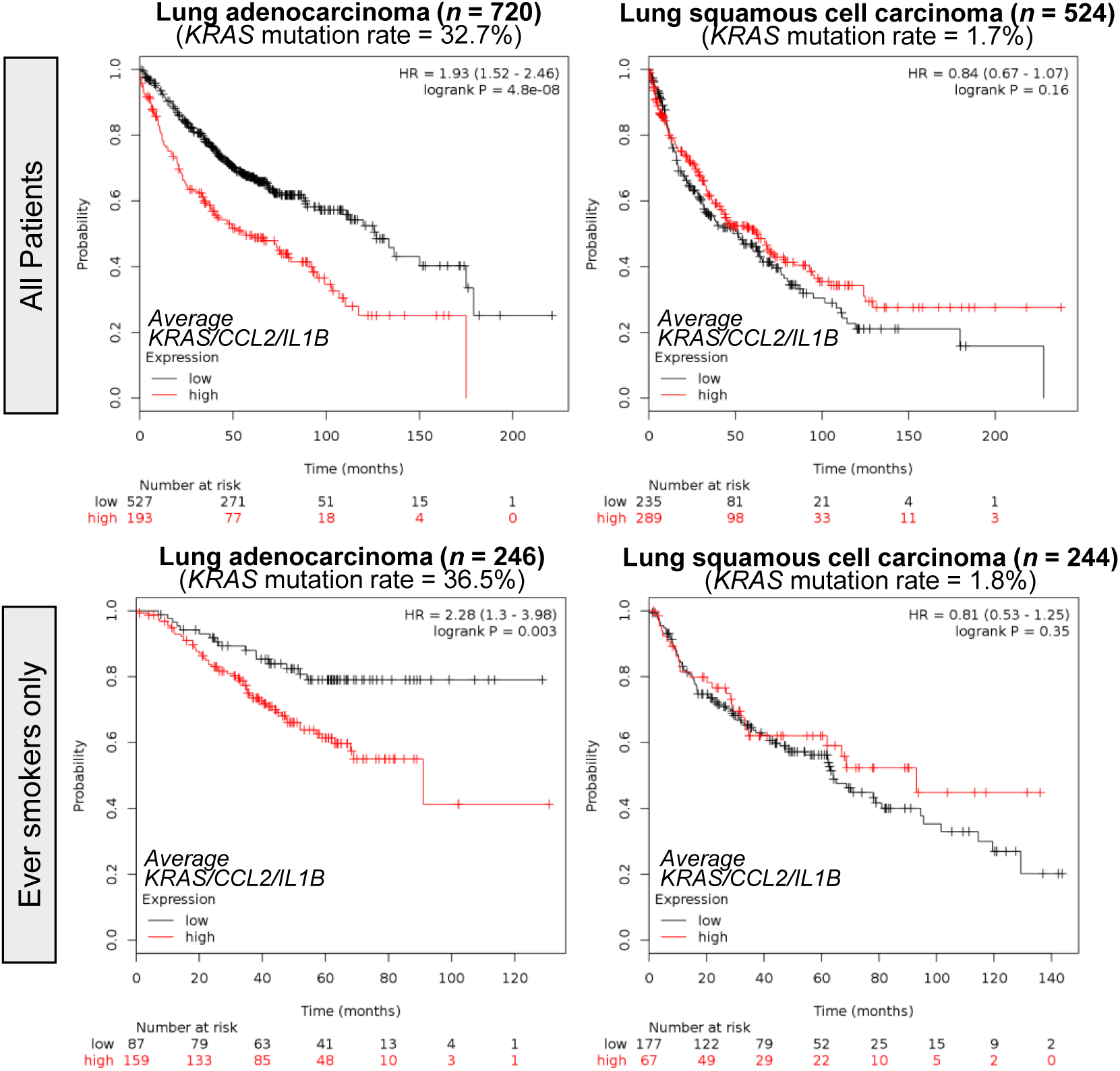
*KRAS/CCL2/IL1B* expression predicts poor survival of *KRAS*-mutant cancers. Kaplan-Meier analyses of lung cancer patients stratified by average *KRAS/CCL2/IL1B* expression done on http://www.kmplot.com. *KRAS* mutation frequencies are from the Campbell cohort (***Campbell et al., 2016***).Top: all patients; Bottom: ever-smokers only.

## DISCUSSION

We hypothesized that mutant *KRAS* dependence occurs non-cell-autonomously and that KRAS inhibitor effects are delivered *in vivo*. We used 30 cancer cell lines with different *KRAS* mutations and multiple *in vitro* assays to show that both pharmacologic and genetic KRAS inhibition is selectively effective against *KRAS*-mutant murine and human tumors *in vivo*. Using isogenic cell lines with intact or compromised mutant *KRAS* signaling, we identify a novel *KRAS*-mutation-specific transcriptome signature that is surprisingly predominated by inflammatory response genes including *CCL2* and *IL1R1*. We further employ several transgenic mouse strains and adoptive bone marrow transfer experiments to show that effective pharmacologic KRAS blockade *in vivo* is dependent on the presence of CCR2+ IL-1β myeloid cells in the tumor microenvironment. Finally, we show that the KRAS blocker deltarasin functions to downregulate *IL1R1* expression in *KRAS*-mutant tumor cells and that the proposed *KRAS/CCL2/IL1B* signature is enriched in human cancers with high *KRAS* mutation frequencies in which it portends a dismal prognosis. Our results imply that conventional cell-based screens for the discovery and development of novel KRAS blockers might be suboptimal, and that IL-1β inhibition may be specifically effective against KRAS-mutant cancers.

A long line of evidence supports that homotypic two-dimensional cancer cell cultures are not optimal for the study of KRAS-dependence. Singh et al. established a “RAS-dependency index” in a large panel of human lung and pancreatic cancer cell lines systematically addressing the variable *in vitro* efficacy of KRAS inhibition (***Singh et al., 2009***). Project DRIVE, a comprehensive synthetic lethality screen applying> 150,000 shRNAs on 7,837 genes and 398 cancer cell lines (https://oncologynibr.shinyapps.io/drive/), identified no lethal interaction partners for KRAS *in vitro*, a finding that urged the authors to state: “… the data here raise the likelihood that no single synthetic lethal gene will be found across all KRAS mutant tumors … commonly used KRAS mutant models are not KRAS dependent, when interrogated as monolayer cell cultures … ablating KRAS dependence will need to carefully consider these findings…” (***McDonald et al., 2017***). Recently, Janes et al. developed ARS-1620, a new covalent G12C-specific KRAS inhibitor that is highly effective *in vivo*, but not *in vitro* (***Janes et al., 2018***). The authors developed three-dimensional co-culture systems and state: “We use ARS-1620 to dissect oncogenic KRAS dependency and demonstrate that monolayer culture formats significantly underestimate KRAS dependency *in vivo*”. Despite the tremendous progress contributed by the above-referenced work, the mechanism(s) of the observed *in vivo*-restricted KRAS-dependence remained obscure prior to this report.

To this end, multiple lines of work support the notion that the paracrine effects of KRAS and other RAS oncogenes overshadow their cell-autonomous impact. A pioneering report identified how RAS oncogenes utilize paracrine IL-8 signaling to induce angiogenesis *in vivo* (***Sparmann and Bar-Sagi, 2004***; ***Karin, 2004***). We determined how *KRAS*-mutant cancer cells depend on paracrine CCL2 signaling to myeloid cells including mononuclear and mast cells to induce vascular permeability and angiogenesis during malignant pleural effusion development (***Giannou et al., 2015***; ***Agalioti et al., 2017***). In turn, myeloid-derived IL-1β was found to selectively trigger non-canonical nuclear factor (NF)-κΒ activation in *KRAS*-mutant cancer cells via IL1R1 and inhibitor of NF-κΒ), with the latter presenting a marked therapeutic target in mouse models of pre-metastatic and advanced lung cancer (***Marazioti et al., 2018***; ***Vreka et al., 2018***). Here we show how deltarasin functions to abrogate a mutant *KRAS*-initiated *in vivo* inflammatory loop of tumor-derived CCL2 and myeloid-secreted IL-1β by downregulating IL1R1 expression of *KRAS*-mutant tumor cells and thereby abolishing their receptivity to myeloid IL-1β signals. We identify CCR2+ myeloid cells that provide IL-1β to the microenvironment of *KRAS*-mutant tumors and show that they are required for mutant *KRAS* dependence *in vivo*. Data from syngeneic mouse models of global host *Ccr2* and *Il1b* gene deficiency and of focal myeloid *Ccr2* reconstitution are further supported by human cancer xenograft experiments in *Rag2^-/-^*mice, which lack B- and T-cell function but feature intact myeloid cells (***Hao and Rajewsky, 2001***), to collectively identify the proposed inflammatory loop that potentiates KRAS blockade.

In addition to *Kras*, *Ccl2*, and *Il1r1*, a battery of other transcripts emanated within the signature of *KRAS*-mutant cancers derived from the transcriptomes of our cell lines, providing synthetic lethality candidates for *in vivo* KRAS dependency for future research. This signature includes signal transducers *Ranbp3l*, *Gpr149*, and *Rassf8*, inflammatory messengers *Ccl7*, *Cxcl1*, and *Casp3*, cell surface receptors *Pdgfra* and *Ttk*, cell cycle genes and tumor suppressors *Cdca5*, *Hist2h3c2*, *Plag1*, *Fanca*, and *Gmnn*, among others. The importance of some of these candidates is worth mentioning: *Cxcl1* was recently found to mediate the effects of KRAS-IKKα addiction during malignant pleural effusion development (***Marazioti et al., 2018***); *Casp3* is a central effector of compensatory tumor proliferation and radiotherapy resistance (***Huang et al., 2011***); and *Gmnn* was recently found to function as a tumor suppressor in lung and colon cancer (***Champeris Tsaniras et al., 2018***). Surprisingly, *Kras* mutation status imprinted the transcriptomes of our cell lines more profoundly than their tissues of origin, making them cluster together in an unsupervised fashion. Furthermore, our *KRAS*-mutation signature was enriched in human *KRAS*-mutant cancers and predicted poor survival, a fact that is further validating this gene set. Most importantly, the mutant *KRAS* signature was dominated by the inflammatory response pathway on both WikiPathway analysis and GSEA, highlighting the notion that the oncogene functions in a proinflammatory fashion.

In addition to fostering the battle to drug KRAS, the present work bears significant clinical implications by pinning CCL2 and IL-1β as key inflammatory addiction partners of mutant *KRAS*. Although targeting CCL2 with neutralizing antibodies yielded promising preclinical results (***Loberg et al., 2007***; ***Fridlender et al., 2010***; ***Qian et al., 2011***; ***Giannou et al., 2015***; ***Agalioti et al., 2017***; ***Marazioti et al., 2013***), clinical trials of the anti-human CCL2 antibody carlumab were hampered by limited drug efficacy and tolerability (***Brana et al., 2015***; ***Sandhu et al., 2013***; ***Pienta et al., 2013***). In contrast, targeting IL-1β with canakinumab has raised enthusiasm and holds great promise in cancer therapy. In this regard, the Canakinumab Anti-inflammatory Thrombosis Outcomes Study (CANTOS), a controlled randomised trial of the role of IL-1β inhibition in atherosclerosis, secondarily aimed at establishing whether low (50 mg), medium (150 mg), or high (300 mg)-dose canakinumab given sc every three months might alter cancer incidence (***Ridker et al., 2017a and 2017b***). The results astonished, with total cancer mortality decreasing by 51% in the high-dose group, incident lung cancer decreasing by 39% in the medium-dose and by 67% in the high-dose groups, and with lung cancer mortality decreasing by 77% in the high-dose canakinumab group. Although our results of diminished deltarasin efficacy with *Il1b^-/-^* mice were less impressive compared with the complete abrogation of deltarasin effects in *Ccl2^-/-^* mice, we believe that this is attributable to redundant IL-1α signaling in the former and that targeting IL-1β might be specifically effective against *KRAS*-mutant cancers (***Song et al., 2003***; ***Voronov et al., 2003***; ***Voigt et al., 2017***; ***Apte and Voronov, 2008***; ***Dinarello et al., 2012***). This is plausible from CANTOS results, since canakinumab effects in decreasing lung cancer incidence and mortality were double in current than in past smokers overall and quadruple when the high-dose group was examined alone, with current smokers having higher *KRAS* mutation rates than never-smokers (***Tate et al., 2019***; ***Campbell et al., 2016***). Our results suggest that canakinumab might be selectively effective against *KRAS*-mutant cancers and warrant *a posteriori* analysis of CANTOS results by *KRAS* mutation status.

In summary, we show that *KRAS*-mutant cancer cells express CCL2 and IL1R1 to initiate an β-expressing myeloid cells. Our work indicates that this crosstalk is required for KRAS-dependence and blockade, which targets IL1R1 expression. The data set a rational framework for the future development of effective KRAS inhibitors and design of clinical trials aimed at targeting IL-1β in cancer.

## MATERIALS AND METHODS

### Key Resources Table

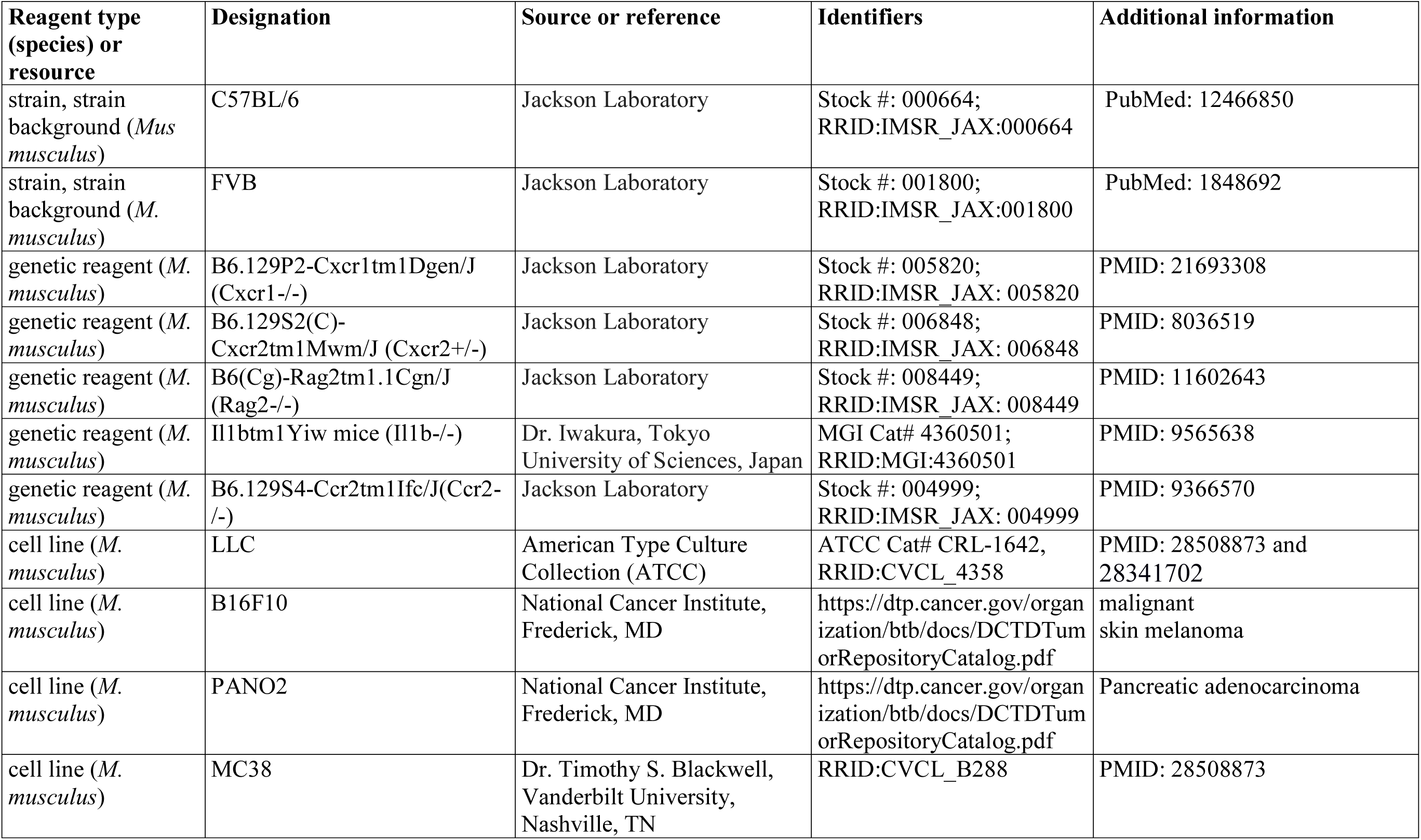
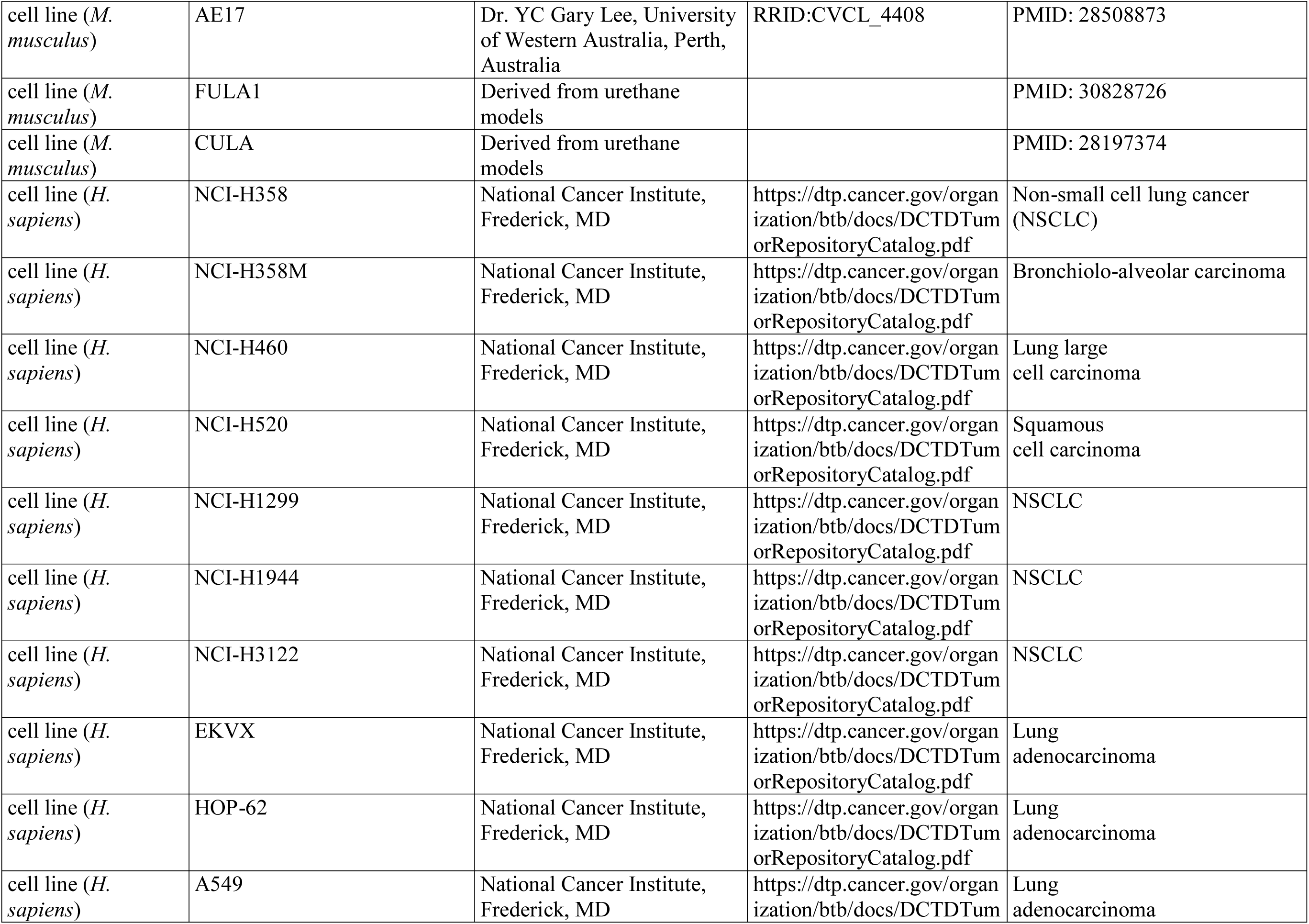
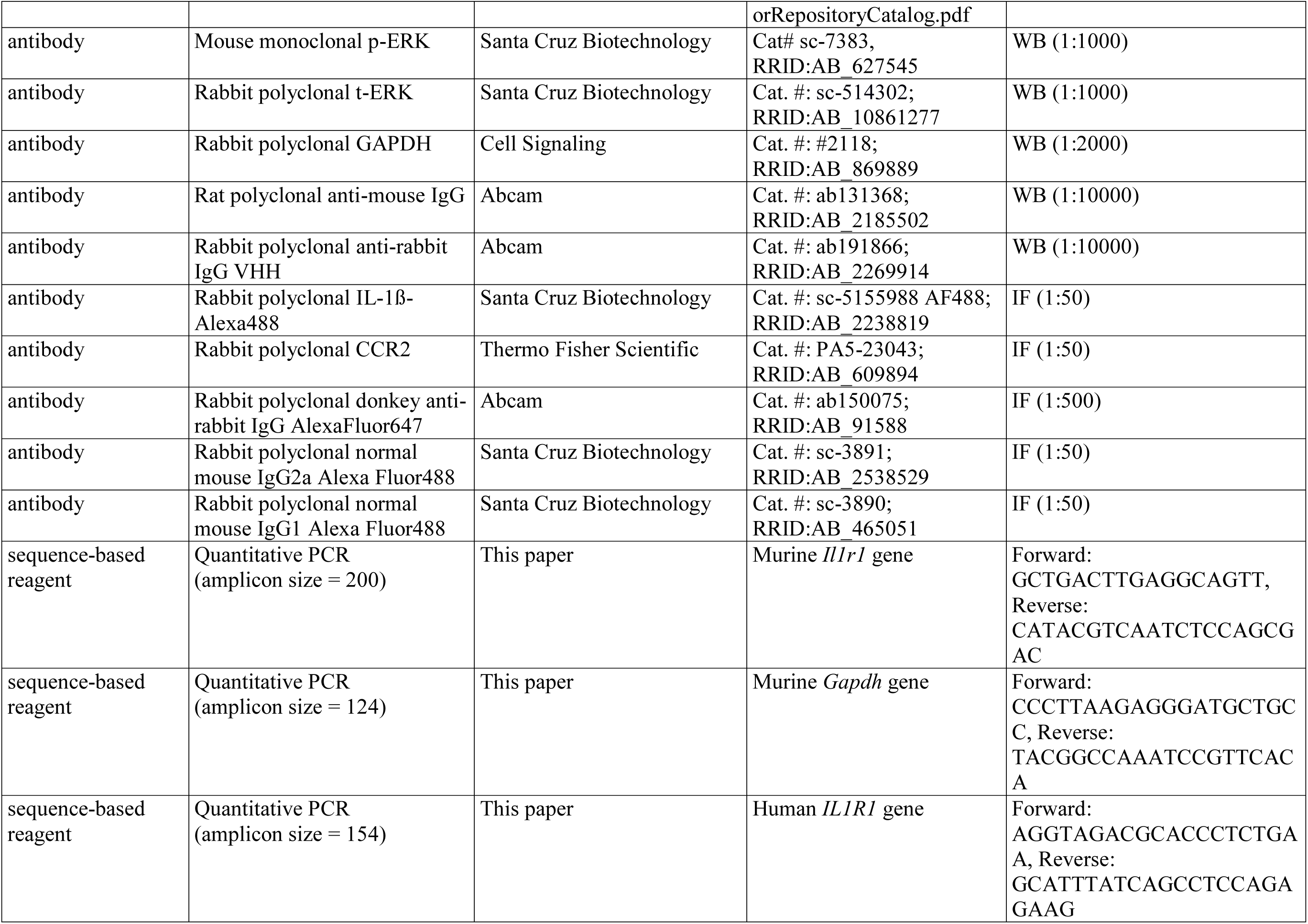
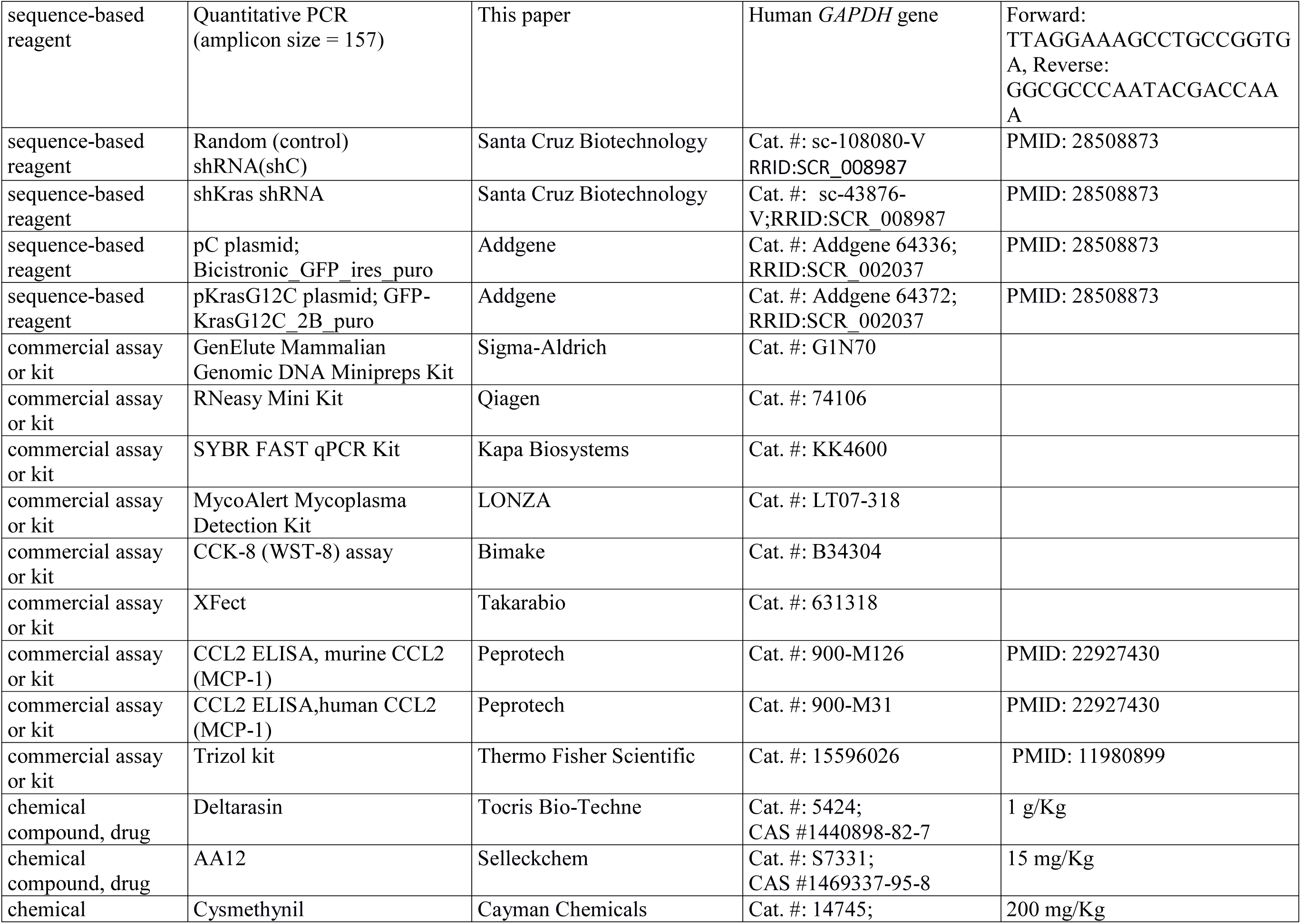
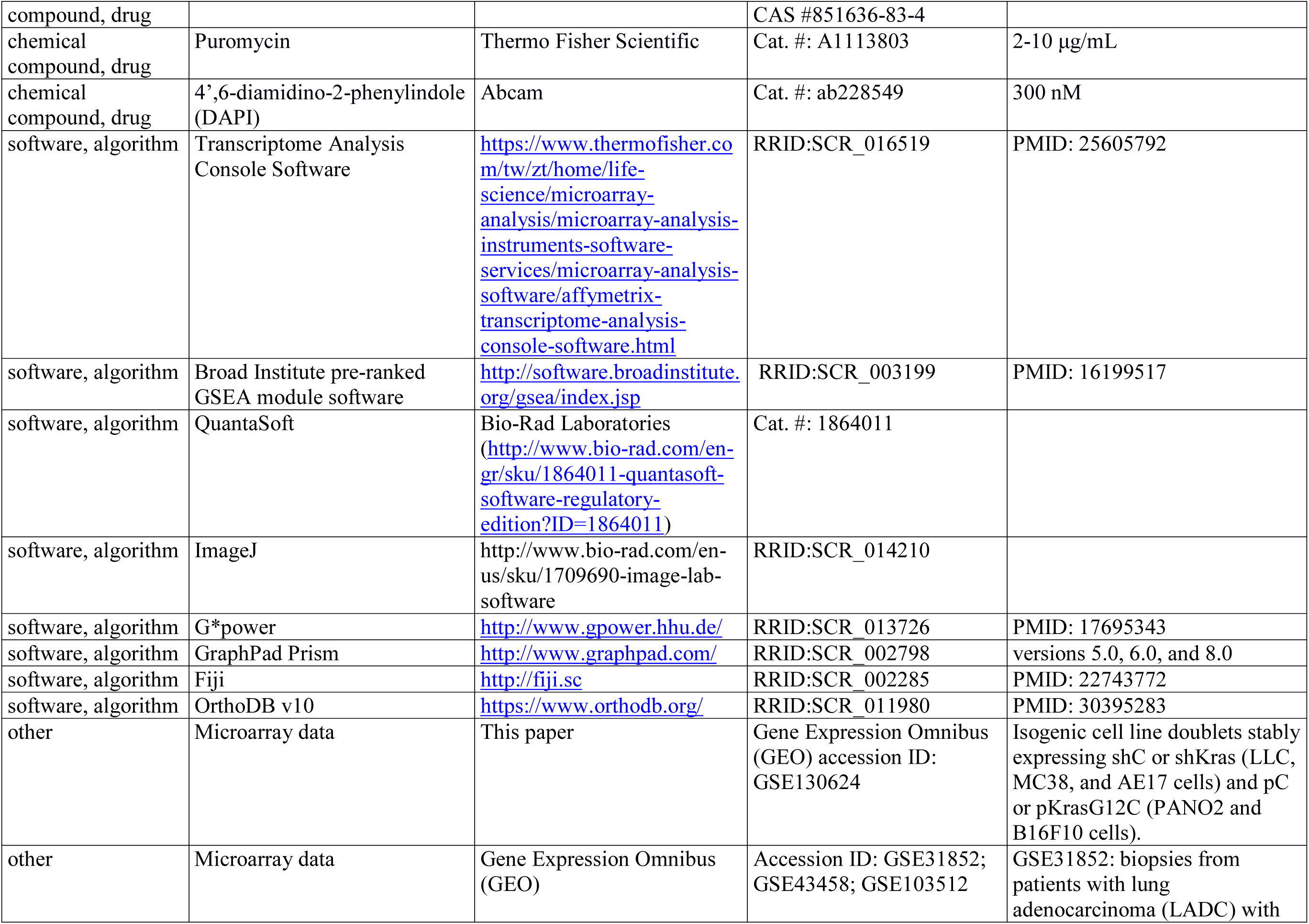
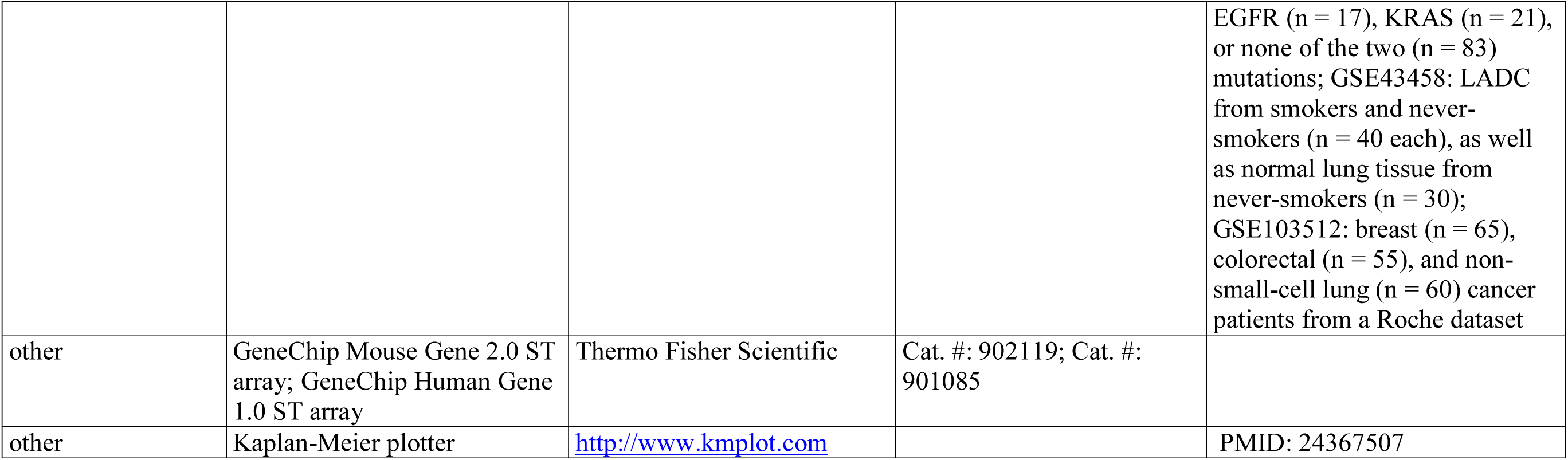

### Cell culture

NCI-H358, NCI-H358M, NCI-H460, NCI-H520, NCI-H1299, NCI-H1944, NCI-H3122 (referred to hereafter omitting NCI), EKVX, A549, LLC, B16F10, and PANO2 cell lines were from the National Cancer Institute (Frederick, MD); MC38 cells were a gift from Dr. Timothy S. Blackwell (Vanderbilt University, Nashville, TN) and AE17 cells from Dr. YC Gary Lee (University of Western Australia, Perth, Australia) (***Agalioti et al., 2017***; ***Marazioti et al., 2018***; ***Giannou et al., 2017***). FULA1 (*FVB* urethane-induced lung adenocarcinoma 1) and CULA (*C57BL/6* urethane-induced lung adenocarcinoma) cell lines were isolated from the lungs of *FVB* and *C57BL/6* mice, respectively, harboring primary lung adenocarcinomas induced by urethane (***Giopanou et al., 2016***; ***Agalioti et al., 2017***; ***Kanellakis et al., 2019***). Human and murine cell lines were cultured, respectively, in Roswell Park Memorial Institute (RPMI)-1640 Dulbecco’s Modified Eagle Medium (DMEM), both supplemented with 10% FBS and 100 and IU/mL penicillin/streptomycin, and were maintained in a humidified incubator at 37 °C with 95% air–5% CO_2_. Cell lines were authenticated annually using the short tandem repeat method and were tested negative for *Mycoplasma Spp.* biannually by MycoAlert Mycoplasma Detection Kit (LONZA; Verviers, Belgium).

### Drugs

Deltarasin (CAS #1440898-82-7; Tocris Bio-Techne #5424; Wiesbaden-Nordenstadt, Germany), KRAS^G12C^ inhibitor 12 (AA12; CAS #1469337-95-8; Selleckchem #S7331; Houston, TX), and cysmethynil (CAS #851636-83-4; Cayman Chemicals #14745; Ann Arbor, MI) were dissolved in DMSO to 10 mM stock concentration and stored at −80 °C. For *in vitro* and *in vivo* experiments, drugs were further diluted in normal saline (resulting in 2% DMSO solutions) and equimolar DMSO solutions (2%) were used as control.

### Cellular Assays

*In vitro* cell proliferation was determined using WST-8 [water soluble tetrazolium-8 or 2-(4-iodophenyl)-3-(4-nitrophenyl)-5-(2,4-disulphophenyl)-2H-teterazolium] assay (Bimake; Munich, Germany). For this, 3000 cells/well were plated in triplicates in 96-well plates in 5% FBS-containing media and allowed to adhere overnight, followed by treatment with different drug concentrations. WST-8 reagent was added 72 hours later according to the manufacturer’s protocol and absorbance at 450 nm was measured 1-4 hours later on a TECAN Sunrise microplate reader (Männedorf, Switzerland). For colony formation assay, 300 cells were plated in triplicates in 6-well plates in 5% FBS-containing media, were treated 24 hours later with 1-2 µM deltarasin, media were replaced with drug-free media 72 hours later, and cells were incubated until ≤ 50 colonies formed. Colonies were fixed with 80% ethanol, stained with 0.5% crystal violet, counted and photographed. All cellular experiments were independently repeated at least thrice.

### Western Immunoblotting

Cellular protein lysates were prepared using radioimmunoprecipitation assay (RIPA) buffer containing phosphatase/protease inhibitor cocktail (Thermo Fisher, Waltham, MA), separated by SDS-PAGE, and transferred to nitrocellulose membranes according to standard protocols. Anti-t-ERK, anti-p-ERK, and anti-GAPDH antibodies were from Santa Cruz Biotechnology (Houston, TX). Blots were developed on a Chemidoc XRS+ System (Biorad Labaratories Inc., Hercules, CA).

### Constructs

Short-hairpin (sh) RNA-mediated *Kras*-silenced (sh*Kras*) LLC, AE17, and MC38 cells, as well as B16F10 and PANO2 cells overexpressing custom-made plasmid encoding *Kras*^G12C^(p*Kras*^G12C^; Addgene #64372; GFP-KrasG12C_2B_puro) were produced as described elsewhere (***Agalioti et al., 2017***). NCI-H3122 and EKVX cells were stably transfected with p*Kras*^G12C^ or its homologous GFP backbone plasmid without *Kras*^G12C^ (p*C*; Addgene #64336; Bicistronic_GFP_ires_puro) using previously established methods (***Agalioti et al., 2017***). All plasmids were made in-house, deposited, validated, and re-purchased from Addgene (Watertown, MA). For stable shRNA and plasmid transfection, 10^5^ tumor cells in 6-well culture vessels were transfected with 5 μg DNA using XFect (Takara, Kusatsu, Japan) and clones were selected by puromycin (2-10 μg/mL).

### Mice

FVB/NJ (*FVB*; #001800), C57BL/6J (*C57BL/6*; #000664), B6.129P2-Cxcr1^tm1Dgen/J^ (*Cxcr1^-/-^*; #005820) (***Sakai et al., 2011***), B6.129S4-Ccr2^tm1Ifc/J^(*Ccr2^-/-^*; #004999) (***Boring et al., 1997***), B6.129S2(C)-*Cxcr2*^tm1Mwm/J^ (*Cxcr2^+/-^*; #006848) (***Cacalano et al., 1994***), and B6(Cg)- *Rag2*^tm1.1Cgn^/J (*Rag2^-/-^*; #008449) (***Hao et al., 2001***) mice were from the Jackson Laboratory (Bar Harbor, ME) and *Il1b*^tm1Yiw^ mice (*Il1b^-/-^*; MGI #2157396) (***Horai et al., 1998***) were a kind gift from Dr. Yoichiro Iwakura (Tokyo University of Sciences, Japan). All mice were bred at the Center for Animal Models of Disease of the University of Patras. *Ccr2^-/-^* mice were back-crossed to the *FVB* strain for > F12. Experimental mice were weight- (20-30 g), sex-, and age- (6-12 weeks) matched; both female and male mice were used and were allocated to treatment groups by alternation. In these studies, 284 mice were enrolled in total. In more detail, 25 *FVB* (21 for tumor experiments and 4 as bone marrow donors), 151 *C57BL/6* (all for tumor experiments), 15 *Cxcr1^-/-^* (all on the *C57BL/6* background for tumor experiments), 34 *Ccr2^-/-^* (12 on the *C57BL/6* and 18 on the *FVB* backgrounds for tumor experiments and 4 on the *FVB* background as bone marrow donors), 12 *Cxcr2^+/-^* (all on the *C57BL/6* background for tumor experiments), 32 *Rag2^-/-^* (all on the *C57BL/6* background for tumor experiments), and 15 *Il1b^-/-^* (all on the *C57BL/6* background for tumor experiments) mice were used.

### *In vivo* tumor models and drug treatments

For *in vivo* injections, 10^6^ cells suspended in 50 µL PBS were implanted subcutaneously (sc) in the rear flank. Tumor dimensions (length, L; width, W; depth, D) were monitored serially using calipers and tumor volume (V) was calculated as *V* = π * *L* * *W* * *D*/6 . Drug treatments were initiated when tumors reached both 100 mm^3^ volume and 14 days latency post-tumor cell injection and consisted of daily intraperitoneal (ip) deltarasin (15 mg/Kg in 100 μL saline 2% DMSO) or 100μL saline 2% DMSO. Animals were monitored daily for sickness and were euthanized using CO_2_ when in distress or when tumors reached 2-3 cm^3^ volume, whichever occurred first.

### Microarrays, PCR, GSEA,and Kaplan-Meier analyses

Isogenic cell line doublets stably expressing sh*C* or sh*Kras* (LLC, MC38, and AE17 cells) and p*C* or p*Kras*^G12C^ (PANO2 and B16F10 cells) were generated as described elsewhere (***Agalioti et al., 2017***). Benign samples including whole murine lungs, tracheal epithelial cells (TEC; cultured out from murine tracheas), and bone marrow-derived macrophages (BMDM; cultured from murine bone marrow by weekly incubation with 20 ng/mL M-CSF) and mast cells (BMMC; cultured from murine bone marrow by monthly incubation with 100 ng/mL IL-3 plus KITL) were prepared as described elsewhere (***Giannou et al., 2015***; ***Marazioti et al., 2018***; ***Kanellakis et al., 2019***; ***Lilis et al., 2019***). Cellular RNA was isolated using Trizol (Thermo Fisher) followed by RNAeasy column purification and genomic DNA removal (Qiagen, Hilden, Germany). One μg RNA was reverse-transcribed using oligo(dT)_18_ and iScript Advanced cDNA synthesis kit for RT-qPCR (Bio-Rad Laboratories; Hercules, CA). *Il1r1/IL1R1* and *Gapdh/GAPDH* qPCR was performed using specific primers and Lightcycler 480 Sybr Green I Master (Roche; Mannheim, Germany) in a Lightcycler 480 II (Roche Diagnostics). Ct values from triplicate reactions were analyzed with the 2^-Δ^ method (***Pfaffl, 2001***) as detailed elsewhere (***Giannou et al., 2017***). mRNA abundance was determined relative to *Gapdh/GAPDH* and is given as 2^-Δ^ = 2*^Il1r1/IL1R1^*^)-(Ct of *Gapdh/GAPDH*)^. Mouse microarrays were done as described elsewhere (***Giannou et al., 2015***; Agalioti et al., 2017; Marazioti et al., 2018; Kanellakis et al., 2019; Lilis et al., 2019). Briefly, triplicate cultures of 10^6^ cells were subjected to RNA extraction as above, 5 μg of pooled total RNA were tested for RNA quality on an ABI2000 Bioanalyzer (Agilent; Santa Clara, CA), labeled, and hybridized to GeneChip Mouse Gene 2.0 ST arrays (Affymetrix; Santa Clara, CA). Analyses using Affymetrix Expression/Transcriptome Analysis Consoles (***Ritchie et al. 2015***) consisted of normalization of all arrays together using Lowess multi-array algorithm, intensity-dependent estimation of noise for statistical analysis of differential expression, and unsupervised hierarchical clustering of microarray data and WikiPathway analysis. Murine microarray data generated for this study are publicly available at the Gene Expression Omnibus (GEO) database (http://www.ncbi.nlm.nih.gov/geo/; Accession ID: GSE130624). Gene set enrichment analyses (GSEA) was done using publicly available Human Gene 1.0 ST microarray data obtained from GEO. The following datasets were used: GSE31852 with gene expression profiles of 121 biopsies from patients with lung adenocarcinoma (LADC) with *EGFR* (*n* = 17), *KRAS* (*n* = 21), or none of the two (*n* = 83) mutations [Biomarker-integrated Approaches of Targeted Therapy for Lung Cancer Elimination (BATTLE) trial]; GSE43458 with gene expression profiles of LADC from smokers and never-smokers (*n* = 40 each), as well as normal lung tissue from never-smokers (*n* = 30) also from the BATTLE trial; and GSE103512 with gene expression profiles of breast (*n* = 65), colorectal (*n* = 55), and non-small-cell lung (*n* = 60) cancer patients from a Roche dataset.Kaplan-Meier analyses were done using KM-plotter (http://www.kmplot.com) (***Gy rffy et al., 2013***). All patients were included and overall survival and all stages/grades were set as parameters.

### ELISA

Murine and human CCL2 levels of cell culture supernatants were detected using appropriate ELISA kits (Peprotech; London, UK). For sample preparation, cells were incubated with IC_60_ deltarasin for 72 hours before collecting cell-free supernatants for CCL2 measurements and whole cellular lysates for normalization of CCL2 levels to total cellular protein.

### Immunofluorescence

Paraffin-embedded mouse tissue blocks were cut into 3 µm-thick sections, deparaffinized by ethanol gradient, rehydrated, and boiled in sodium citrate pH 6.0 for 10 minutes for antigen retrieval. After post-fixation and permeabilization, tissue sections were co-stained with either AlexaFluor488-conjugated mouse monoclonal anti-IL-1β antibody and rabbit polyclonal anti-CCR2 antibody or AlexaFluor488-conjugated mouse monoclonal anti-IL1R1 antibody and rabbit polyclonal anti-CCL2 antibody. After counterstaining with 300 nM 4’,6-diamidino-2-phenylindole (DAPI), slides were evaluated on an AxioImager.M2 (Zeiss; Jena, Germany) and digital images were processed with Fiji academic software (https://fiji.sc/) (***Schindelin et al., 2012***). Control stains were carried out with isotype controls for normal mouse IgG1/ IgG2a (Alexa Fluor® 488 conjugated; sc-3891/ sc-3890) and secondary antibody only.

### Bone marrow replacement

For adoptive bone marrow transplants (BMT), bone marrow cells were flushed from both femurs and tibias of wild-type (*WT*) or *Ccr2^-/-^* mice (all back-crossed >F12 to the *FVB* background) using fully supplemented DMEM. *Ccr2^-/-^* mice (all *FVB*) received ten million intravenous (iv) bone marrow cells from *WT* or *Ccr2^-/-^* mice 12 hours after total-body irradiation (900 Rad), as described elsewhere (***Giannou et al., 2015***; ***Marazioti et al., 2018***). One mouse in each experiment was not engrafted and was observed till moribund on days 5-15 post-irradiation. One month was allowed for full bone marrow reconstitution of chimeras prior to tumor cell injections, as described and validated previously (***Giannou et al., 2015***; ***Marazioti et al., 2018***).

### Statistics

Sample size was calculated using G*power (http://www.gpower.hhu.de/) (***Faul et al., 2007***). In specific, we set out to determine biologically (> 50%) and statistically (α = 0.05; β= 0.20) significant differences between two unmatched independent groups with SD ∼ 30% of mean using two-tailed t-tests, yielding *n* = 7/group. Hence experiments with *n* = 5 mice/group were contemplated in batches, till the achievement of probability (*P*) < 0.05 with α < 0.05 or *P*> 0.05 with β < 0.20, whichever came first. All *in vitro* experiments were performed at least three independent times (biological replicates), each time using at least three technical replicates. All source data are provided as *.xlsx source data files and all data were included in the analyses without elimination of outliers. Two-way ANOVA was employed to achieve further reduction. Results are given as mean ± SD. Sample size (*n*) refers to biological replicates. Differences between means were assessed using one-way or two-way ANOVA with Bonferroni post-tests. Fifty and 60% inhibitory concentrations (IC_50/60_) were calculated using nonlinear regression, a logarithmic inhibitor-response model, unweighted least squares regression without outlier elimination and constraints, and extra sum-of-squares F-test comparisons. *P*< 0.05 was considered significant. Statistics and plots were done on Prism versions 5.0, 6.0, and 8.0 (GraphPad; San Diego, CA).

### Study approval

Experiments were approved by the Veterinary Administration of the Prefecture of Western Greece (approval # 366456/1461) and by the Government of Upper Bavaria (approval # 55.2-1-54-2532-194-2016) and were conducted according to Directive 2010/63/EU (http://eurlex.europa.eu/legal-content/EN/TXT/?uri=CELEX%3A32010L0063).

## DATA AVAILABILITY

All raw data produced in this study are provided as *.xlsx source data supplements. The microarray data produced by this study were deposited at GEO (http://www.ncbi.nlm.nih.gov/geo/; Accession ID: GSE130624). Gene set enrichment analyses (GSEA) were done using publicly available microarray data obtained from GEO (http://www.ncbi.nlm.nih.gov/geo/; Accession IDs: GSE31852, GSE43458, and GSE103512).

## Supporting information

Title page metadata

## LEGENDS TO FIGURES, FIGURE SUPPLEMENTS& SOURCE DATA

**Figure 1–figure supplement 1.**
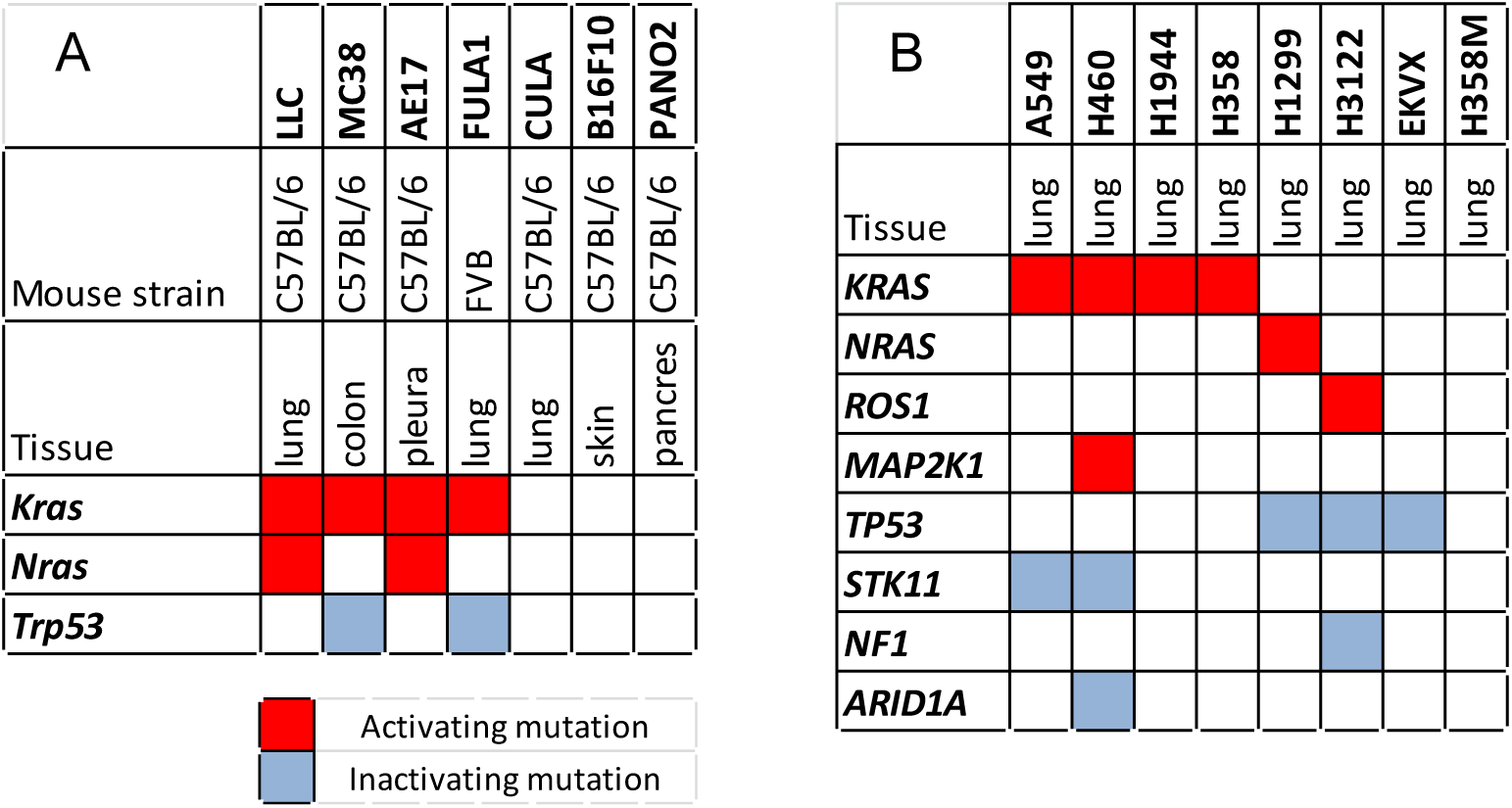
Mutation status of cell lines used in this study. Cell lines used in this study with their syngeneic mouse strain, tissue of origin, and mutation status. Data from Giopanou et al., 2016; Agalioti et al., 2017; Giannou et al., 2017; Marazioti et al., 2018;Tate et ***al., 201****9*; and ***Kanellakis et al., 2019***.

**Figure 1–figure supplement 2.**
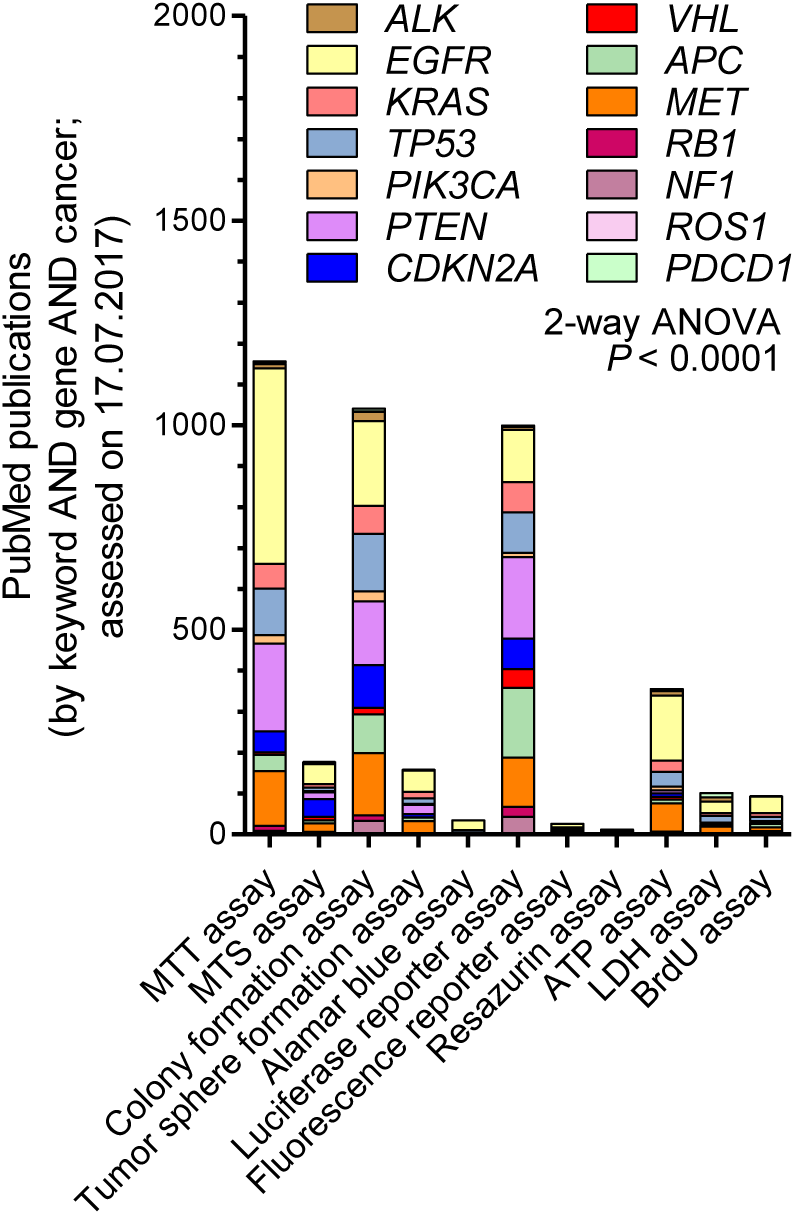
*In vitro* assays used in cancer research. Summary of *in vitro* assays used in cancer research stratified by target gene. Data are from a PubMed search done between 17-29.07.2018 using search strategy (“assay type” AND “gene” AND “cancer”) and the number of retrieved publications as the readout. Assay types are listed in the x-axis and genes in the legend.MTT,3-(4,5-dimethylthiazol-2-yl)-2,5-diphenyltetrazolium bromide;MTS,3-(4,5-dimethylthiazol-2-yl)-5-(3-carboxymethoxyphenyl)-2-(4-sulfophenyl)-2H-tetrazolium); ATP, adenosine triphosphate; LDH, Lactate dehydrogenase; BrdU, bromodeoxyuridine, 5-bromo-2’-deoxyuridine. *P*, overall probability by 2-way ANOVA. Note that MTT/MTS and colony formation assays are the most commonly used and were also used in this study.

**Figure 1–figure supplement 3.**
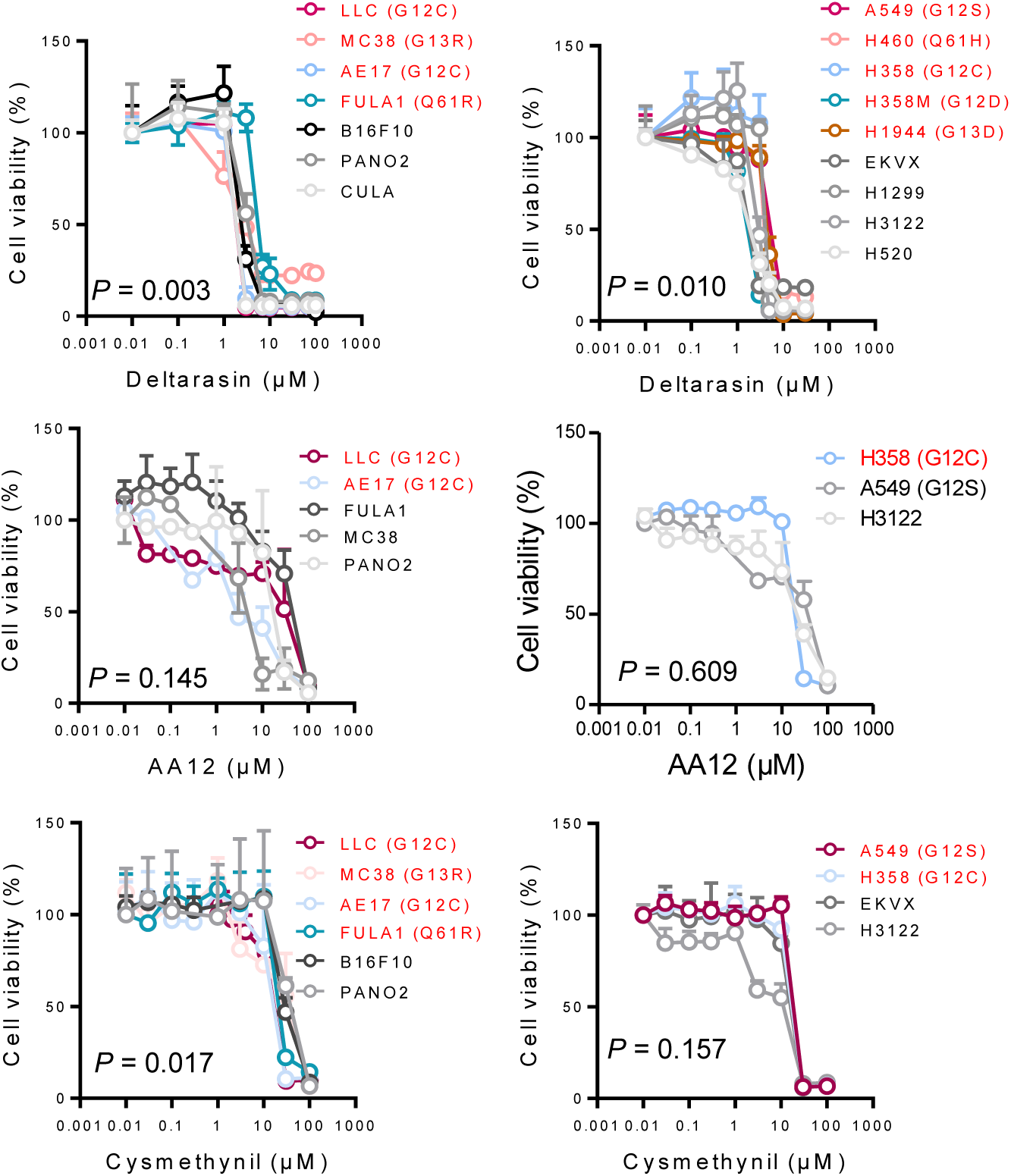
Response of *KRAS*-mutant tumor cells to *KRAS* inhibitors analyzed by WST-8 assay. Different mouse (top; *Kras*^MUT^: LLC, MC38, AE17, FULA1; *Kras*^WT^: B16F10, CULA, PANO2) and human (bottom; *KRAS*^MUT^: A549, H460, H358, H358M, H1944, HOP-62; *KRAS*^WT^: EKVX, H1299, H3122, H520) tumor cell lines were assessed for inhibition of cell viability (determined by WST-8 assay, *n* = 3/data-point) by three different KRAS inhibitors: deltarasin (top), AA12 (middle), and cysmethynil (bottom). Data presented as mean ± SD. *P*, overall probability by nonlinear fit and extra sum of squares F-test.

**Figure 1–figure supplement 4.**
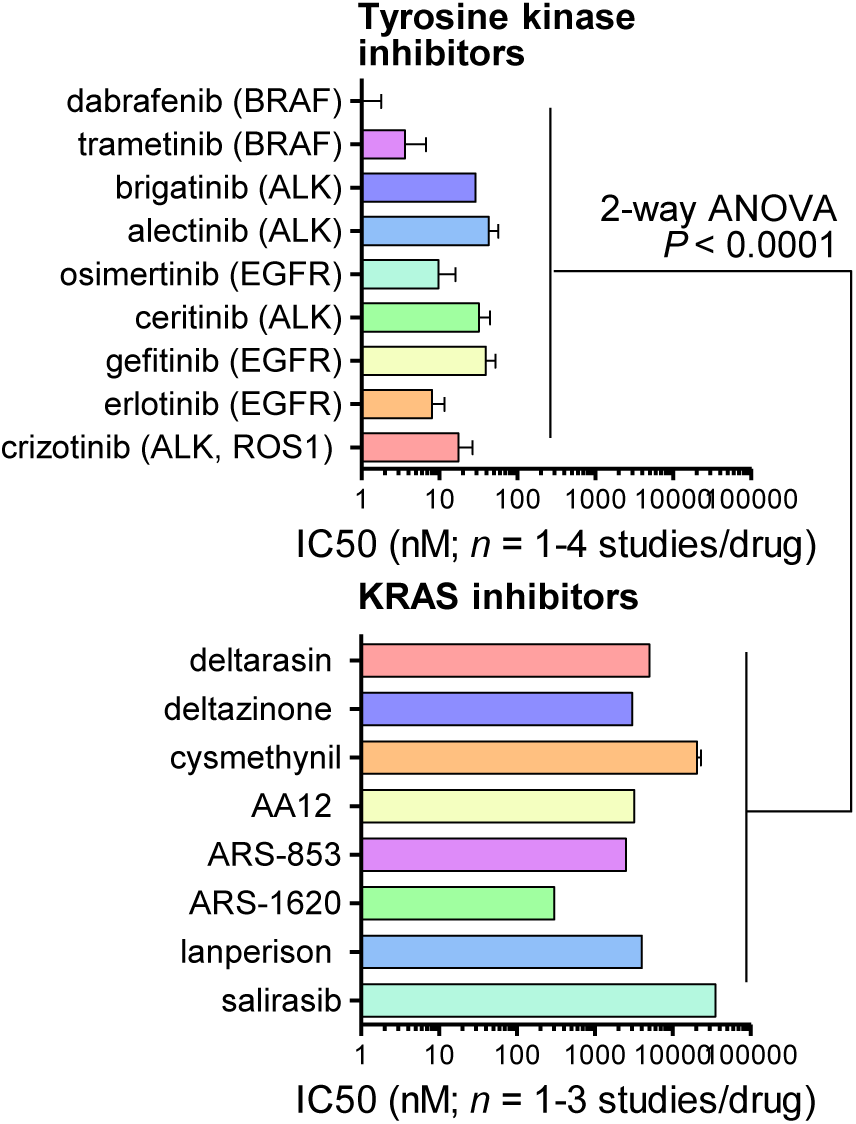
Comparative efficacy of KRAS versus tyrosine kinase inhibitors. Fifty percent inhibitory concentrations (IC_50_) of selected FDA-approved tyrosine kinase inhibitors (TKI; top) and of published KRAS inhibitors(bottom) in preclinical development. *n*, published studies; *P*, overall probability by 2-way ANOVA. Note the statistically significantly higher and physiologically difficult to achieve IC_50_ of KRAS inhibitors compared with TKI. Data were from ***Hong et al., 2011***; ***Yamaguchi et al., 2011***; ***Hirano et al., 2015***;Sakamoto et al., 2011;Huang et al., 2016;Chen et al., 2013;Prahallad et al., 2012;Wilson ***et al., 201****2*;***Zhang et al., 2012***;***Zhang et al., 2016***;***Lito et al., 2016***;***Ostrem et al., 2013***;***Winter– Vann et al., 2005***;***Wang et al., 2010***;***Weisz et al., 1999***;***Zimmermann et al., 2013***;***Shaw et al., 2011***; and ***Papke et al., 2016***.

**Figure 2–figure supplement 1.**
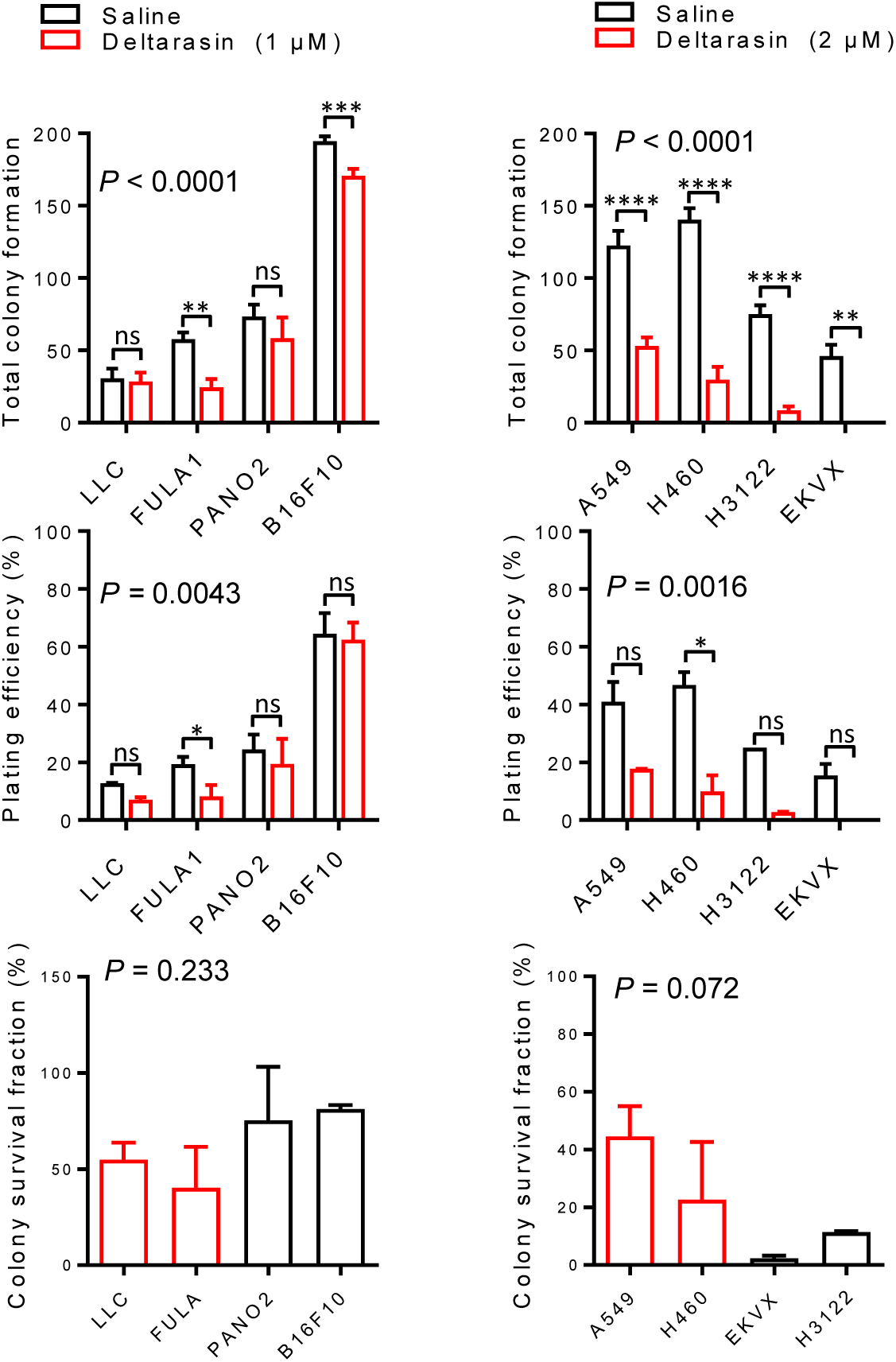
Response of *KRAS*-mutant tumor cells to *KRAS* inhibitors analyzed by colony formation assay. Different mouse (left; *Kras*^MUT^: LLC, FULA1; *Kras*^WT^: B16F10, PANO2) and human (right; *KRAS*^MUT^: A549, H460; *KRAS*^WT^: EKVX, H3122) tumor cell lines were assessed for colony formation (*n* = 3/ data-point) after 72 h of saline or deltarasin treatment. Data presented as mean ± SD. *P*, overall probability by one-way ANOVA. * and ***: *P*< 0.05 and *P*< 0.001, respectively, for the indicated comparisons by Bonferroni post-tests. Shown are total number of colonies formed(top), plating efficiency of 300 cells/well at experiment start (middle), and survival fraction of single cells given as ratio treatment/no treatment.

**Figure 2–figure supplement 2.**
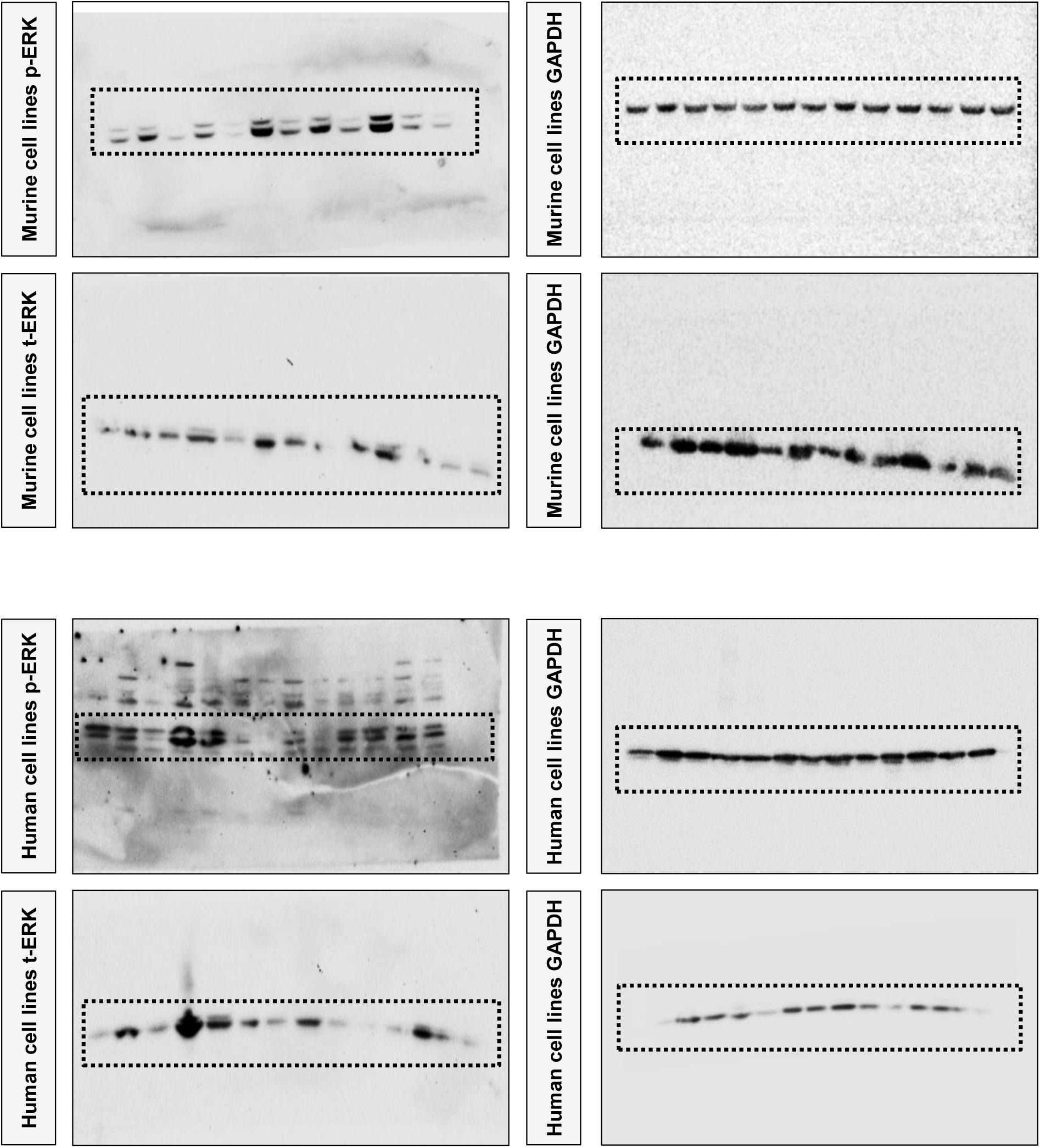
Uncropped blots for Figure 2D. Top: Immunoblots of murine cell line protein extracts untreated and treated with deltarasin (72 h; IC_60_). Left, p-ERK, t-ERK; right, GAPDH. Bottom: Immunoblots of human cell line protein extracts untreated and treated with deltarasin (72 h; IC_60_). Left, p-ERK, t-ERK; right, GAPDH. Dashed lines represent areas of the blots shown in main Figure.

**Figure 3–figure supplement 1.**
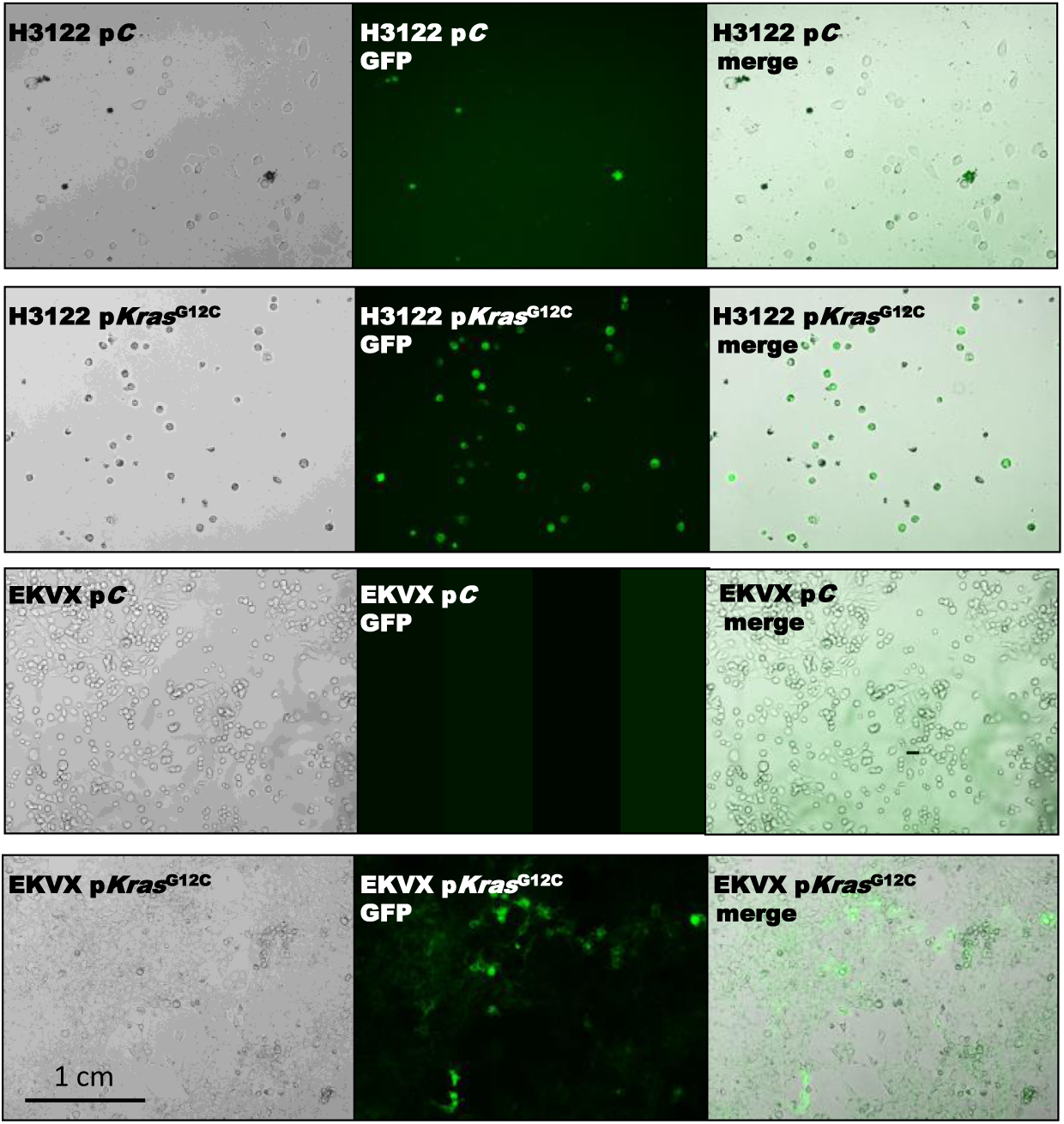
Validation of p*Kras*^G12C^ transduction in human cell lines H3122 and EKVX. Thep*Kras*^G12C^plasmid includes GFP and puromycin resistance genes. Representative microscopy images of p*C* control or p*Kras*^G12C^ transfected cell lines. Left, brightfield images; middle, green fluorescent images; right, merged images. Images were taken with a confocal microscope LCI510 (Zeiss; Jena, Germany).

**Figure 4–figure supplement 1.**
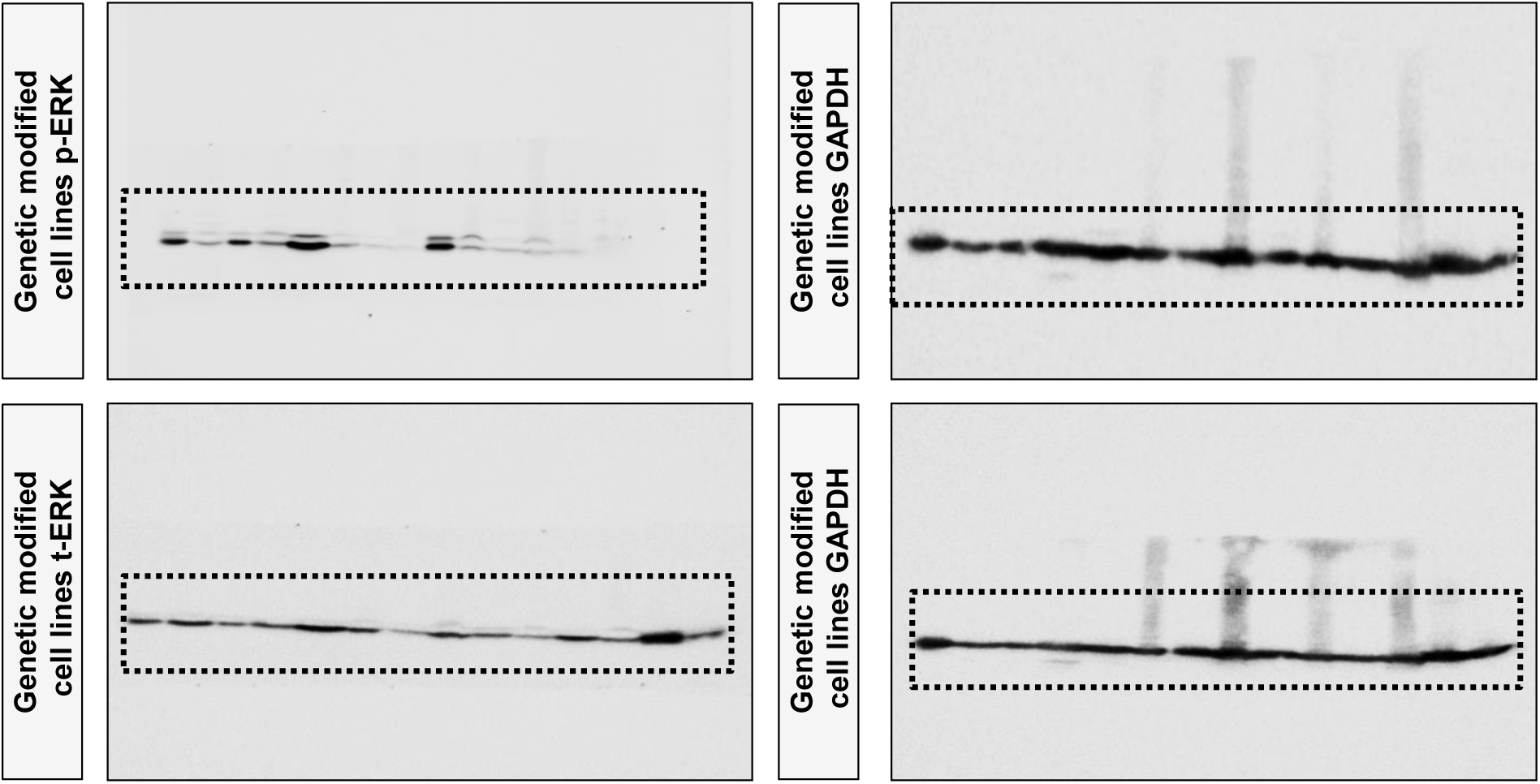
Uncropped blots for Figure 4D. Immunoblots of murine and human cell line protein extracts with or without *Kras/KRAS* genetic modification. Left, p-ERK, t-ERK; right, GAPDH. Dashed lines represent areas of the blots shown in main Figure.

**Figure 1–source data 1.** Source data for Figure 1–figure supplement 2.

**Figure 1–source data 2.** Source data for Figure 1B-D and Figure 1–figure supplement 3.

**Figure 1–source data 3.** Source data for Figure 1–figure supplement 4.

**Figure 2–source data 1.** Source data for Figure 2B and Figure 2–figure supplement 1.

**Figure 2–source data 2.** Source data for Figure 2C.

**Figure 3–source data 1.** Source data for Figure 3A and B.

**Figure 2–source data 2.** Source data for Figure 3C.

**Figure 4–source data 1.**Source data for Figure 4.

**Figure 5–source data 1.** Source data for Figure 5.

**Figure 6–source data 1.** Source data for Figure 6B.

**Figure 6–source data 2.** Source data for Figure 6D.

**Figure 8–source data 1.** Source data for Figure 8C.

**Figure 9–source data 1.** Source data for Figure 9A.

**Figure 9–source data 2.** Source data for Figure 9B.

**Figure 10–source data 1.** Source data for Figure 10A.

**Figure 10–source data 2.** Source data for Figure 10B.

## REFERENCES

Agalioti T, Giannou AD, Krontira AC, Kanellakis NI, Kati D, Vreka M, Pepe M, Spella M, Lilis I, Zazara DE, Nikolouli E, Spiropoulou N, Papadakis A, Papadia K,Voulgaridis A, Harokopos V, Stamou P, Meiners S, Eickelberg O, Snyder LA, Antimisiaris SG, Kardamakis D, Psallidas I, Marazioti A, Stathopoulos GT. 2017. Mutant KRAS promotes malignant pleural effusion formation. Nature Communications 8:15205. DOI: 10.1038/ncomms15205, PMID: 28508873

Apte RN, Voronov E. 2008. Is interleukin-1 a good or bad ’guy’ in tumor immunobiology and immunotherapy? Immunological Reviews222:222–241. DOI: 10.1111/j.1600-065X.2008.00615.x, PMID: 18364005

Boring L, Gosling J, Chensue SW, Kunkel SL, Farese RV Jr, Broxmeyer HE, Charo IF. 1997. Impaired monocyte migration and reduced type 1 (Th1) cytokine responses in C-C chemokine receptor 2 knockout mice. Journal of Clinical Investigation100:2552–2561. DOI: 10.1172/JCI119798, PMID: 9366570

Brana I, Calles A, LoRusso PM, Yee LK, Puchalski TA, Seetharam S, Zhong B, de Boer CJ, Tabernero J, Calvo E. 2015. Carlumab, an anti-C-C chemokine ligand 2 monoclonal antibody, in combination with four chemotherapy regimens for the treatment of patients with solid tumors: an open-label, multicenter phase 1b study. Targets in Oncology10:111–123. DOI: 10.1007/s11523-014-0320-2, PMID: 24928772

Brouwer-Visser J, Cheng WY, Bauer-Mehren A, Maisel D, Lechner K, Andersson E, Dudley JT, Milletti F. 2018. Regulatory T-cell Genes Drive Altered Immune Microenvironment in Adult Solid Cancers and Allow for Immune Contextual Patient Subtyping. Cancer Epidemiology Biomarkers and Prevention 27:103–112. DOI: 10.1158/1055-9965.EPI-17-0461, PMID: 29133367

Cacalano G, Lee J, Kikly K, Ryan AM, Pitts-Meek S, Hultgren B, Wood WI, Moore MW. 1994. Neutrophil and B cell expansion in mice that lack the murine IL-8 receptor homolog. Science265:682–684. DOI: 10.1126/science.8036519, PMID: 8036519

Campbell JD, Alexandrov A, Kim J, Wala J, Berger AH, Pedamallu CS, Shukla SA, Guo G, Brooks AN, Murray BA, Imielinski M, Hu X, Ling S, Akbani R, Rosenberg M, Cibulskis C, Ramachandran A, Collisson EA, Kwiatkowski DJ, Lawrence MS, Weinstein JN, Verhaak RG, Wu CJ, Hammerman PS, Cherniack AD, Getz G; Cancer Genome Atlas Research Network, Artyomov MN, Schreiber R, Govindan R, Meyerson M. 2016. Distinct patterns of somatic genome alterations in lung adenocarcinomas and squamous cell carcinomas. Nature Genetics 48:607–616. DOI: 10.1038/ng.3564, PMID: 27158780

Champeris Tsaniras S, Villiou M, Giannou AD, Nikou S, Petropoulos M, Pateras IS, Tserou P, Karousi F, Lalioti ME, Gorgoulis VG, Patmanidi AL, Stathopoulos GT, Bravou V, Lygerou Z, Taraviras S. 2018. Geminin ablation in vivo enhances tumorigenesis through increased genomic instability. Journal of Pathology246:134–140. DOI: 10.1002/path.5128, PMID: 29952003

Chen J, Jiang C, Wang S. 2013. LDK378: a promising anaplastic lymphoma kinase (ALK) inhibitor. Journal of Medicinal Chemistry 56:5673–5674. DOI: 10.1021/jm401005u, PMID: 23837797

Dinarello CA, Simon A, van der Meer JW. 2012. Treating inflammation by blocking interleukin-1 in a broad spectrum of diseases. Nature Reviews Drug Discovery11:633–652. DOI: 10.1038/nrd3800, PMID: 22850787

Downward J. 2003. Targeting RAS signalling pathways in cancer therapy. Nature Reviews Cancer 3:11–22. DOI: 10.1038/nrc969, PMID: 12509763

Esposito D, Stephen AG, Turbyville TJ, Holderfield M. 2019. New weapons to penetrate the armor: Novel reagents and assays developed at the NCI RAS Initiative to enable discovery of RAS therapeutics. Seminars in Cancer Biology 54:174–182. DOI: 10.1016/j.semcancer.2018.02.006, PMID:29432816

Faul F, Erdfelder E, Lang AG, Buchner A. 2007. G*Power 3: a flexible statistical power analysis program for the social, behavioral, and biomedical sciences. Behavior Research Methods39:175–191. DOI: 10.3758/BF03193146, PMID: 17695343

Fridlender ZG, Buchlis G, Kapoor V, Cheng G, Sun J, Singhal S, Crisanti MC, Wang LC, Heitjan D, Snyder LA, Albelda SM. 2010. CCL2 blockade augments cancer immunotherapy. Cancer Research70:109–118. DOI: 10.1158/0008-5472.CAN-09-2326, PMID: 20028856

Giannou AD, Marazioti A, Kanellakis NI, Giopanou I, Lilis I, Zazara DE, Ntaliarda G, Kati D, Armenis V, Giotopoulou GA, Krontira AC, Lianou M, Agalioti T, Vreka M, Papageorgopoulou M, Fouzas S, Kardamakis D, Psallidas I, Spella M, Stathopoulos GT. 2017. NRAS destines tumor cells to the lungs. EMBO Molecular Medicine 9:672–686. DOI: 10.15252/emmm.201606978, PMID: 28341702

Giannou AD, Marazioti A, Spella M, Kanellakis NI, Apostolopoulou H, Psallidas I, Prijovich ZM, Vreka M, Zazara DE, Lilis I, Papaleonidopoulos V, Kairi CA, Patmanidi AL, Giopanou I, Spiropoulou N, Harokopos V, Aidinis V, Spyratos D, Teliousi S, Papadaki H, Taraviras S, Snyder LA, Eickelberg O, Kardamakis D, Iwakura Y, Feyerabend TB, Rodewald HR, Kalomenidis I, Blackwell TS, Agalioti T, Stathopoulos GT. 2015. Mast cells mediate malignant pleural effusion formation. Journal of Clinical Investigation125:2317–2334. DOI: 10.1172/JCI79840, PMID: 25915587

Giopanou I, Lilis I, Papaleonidopoulos V, Agalioti T, Kanellakis NI, Spiropoulou N, Spella M, Stathopoulos GT. 2016. Tumor-derivedosteopontin isoforms cooperate with TRP53 and CCL2 to promote lung metastasis. Oncoimmunology 6:e1256528. DOI: 10.1080/2162402X.2016.1256528, PMID: 28197374.

Győrffy B, Surowiak P, Budczies J, Lánczky A. 2013. Online survival analysis software to assess the prognostic value of biomarkers using transcriptomic data in non-small-cell lung cancer. PLoS One8:e82241. DOI: 10.1371/journal.pone.0082241, PMID: 24367507

Hao Z, Rajewsky K. 2001. Homeostasis of peripheral B cells in the absence of B cell influx from the bone marrow. Journal of Experimental Medicine 194:1151–1164. DOI: 10.1084/jem.194.8.1151, PMID: 11602643

Hirano T, Yasuda H, Tani T, Hamamoto J, Oashi A, Ishioka K, Arai D, Nukaga S, Miyawaki M, Kawada I, Naoki K, Costa DB, Kobayashi SS, Betsuyaku T, Soejima K. 2015. Invitro modeling to determine mutation specificity of EGFR tyrosine kinase inhibitors against clinically relevant EGFR mutants in non-small-cell lung cancer. Oncotarget 6:38789–803. DOI: 10.18632/oncotarget.5887, PMID: 26515464

Hong S, Hong S, Han SB. 2011. Overcoming metastatic melanoma with BRAF inhibitors. Archives of Pharmaceutical Research 34:699–701. DOI: 10.1007/s12272-011-0521-5, PMID: 21656352

Horai R, Asano M, Sudo K, Kanuka H, Suzuki M, Nishihara M, Takahashi M, Iwakura Y. 1998. Production of mice deficient in genes for interleukin (IL)-1alpha, IL-1beta, IL-1alpha/beta, and IL-1 receptor antagonist shows that IL-1beta is crucial in turpentine-induced fever development and glucocorticoid secretion. Journal of Experimental Medicine 187:1463–1475. DOI: 10.1084/jem.187.9.1463, PMID: 9565638

Huang Q, Li F, Liu X, Li W, Shi W, Liu FF, O’Sullivan B, He Z, Peng Y, Tan AC, Zhou L, Shen J, Han G, Wang XJ, Thorburn J, Thorburn A, Jimeno A, Raben D, Bedford JS, Li CY. 2011. Caspase 3-mediated stimulation of tumor cell repopulation during cancer radiotherapy. Nature Medicine17:860–866. DOI: 10.1038/nm.2385, PMID: 21725296

Huang WS, Liu S, Zou D, Thomas M, Wang Y, Zhou T, Romero J, Kohlmann A, Li F, Qi J, Cai L, Dwight TA, Xu Y, Xu R, Dodd R, Toms A, Parillon L, Lu X, Anjum R, Zhang S, Wang F, Keats J, Wardwell SD, Ning Y, Xu Q, Moran LE, Mohemmad QK, Jang HG, Clackson T, Narasimhan NI, Rivera VM, Zhu X, Dalgarno D, Shakespeare WC. 2016. Discovery of Brigatinib (AP26113), a Phosphine Oxide-Containing, Potent, Orally Active Inhibitor of Anaplastic Lymphoma Kinase. Journal of Medicinal Chemistry 59:4948–64. DOI: 10.1021/acs.jmedchem.6b00306, PMID: 27144831

Janes MR, Zhang J, Li LS, Hansen R, Peters U, Guo X, Chen Y, Babbar A, Firdaus SJ, Darjania L, Feng J, Chen JH, Li S, Li S, Long YO, Thach C, Liu Y, Zarieh A, Ely T, Kucharski JM, Kessler LV, Wu T, Yu K, Wang Y, Yao Y, Deng X, Zarrinkar PP, Brehmer D, Dhanak D, Lorenzi MV, Hu-Lowe D, Patricelli MP, Ren P, Liu Y. 2018. Targeting KRAS Mutant Cancers with a Covalent G12C-Specific Inhibitor. Cell 172:578–589.e17. DOI: 10.1016/j.cell.2018.01.006, PMID:29373830

Kabbout M, Garcia MM, Fujimoto J, Liu DD, Woods D, Chow CW, Mendoza G, Momin AA, James BP, Solis L, Behrens C, Lee JJ, Wistuba II, Kadara H. 2013. ETS2 mediated tumor suppressive function and MET oncogene inhibition in human non-small cell lung cancer. Clinical Cancer Research19:3383–3395. DOI: 10.1158/1078-0432.CCR-13-0341, PMID: 23659968

Kanellakis NI, Giannou AD, Pepe MA, Αgalioti T, Zazara DE, Giopanou I, Psallidas I, Spella M, Μarazioti A, Arendt KAM, Lamort AS, Tsaniras SC, Taraviras S, Papadaki H, Lilis I, Stathopoulos GT. 2019. Tobacco chemical-induced mouse lung adenocarcinoma cell lines pin the prolactin orthologue proliferin as a lung tumour promoter. Carcinogenesis 25:pii:bgz047. DOI:10.1093/carcin/bgz047, PMID: 30828726

Karin M. 2005. Inflammation and cancer: the long reach of Ras. Nature Medicine 11:20–21. DOI: 10.1038/nm0105-20, PMID: 15635437

Kelder T, van Iersel MP, Hanspers K, Kutmon M, Conklin BR, Evelo CT, Pico AR. WikiPathways: building research communities on biological pathways. 2012. Nucleic Acids Research 40:D1301–7. DOI: 10.1093/nar/gkr1074, PMID: 22096230

Kim ES, Herbst RS, Wistuba II, Lee JJ, Blumenschein GR Jr, Tsao A, Stewart DJ, Hicks ME, Erasmus J Jr, Gupta S, Alden CM, Liu S, Tang X, Khuri FR, Tran HT, Johnson BE, Heymach JV, Mao L, Fossella F, Kies MS, Papadimitrakopoulou V, Davis SE, Lippman SM, Hong WK. 2011. The BATTLE trial: personalizing therapy for lung cancer. Cancer Discovery 1:44–53. DOI: 10.1158/2159-8274.CD-10-0010, PMID: 22586319

Kriventseva EV, Kuznetsov D, Tegenfeldt F, Manni M, Dias R, Simão FA, Zdobnov EM. 2019. OrthoDB v10: sampling the diversity of animal, plant, fungal, protist, bacterial and viral genomes for evolutionary and functional annotations of orthologs. Nucleic Acids Research47:D807–D811. DOI: 10.1093/nar/gky1053, PMID: 30395283

Lilis I, Ntaliarda G, Papaleonidopoulos V, Giotopoulou GA, Oplopoiou M, Marazioti A, Spella M, Marwitz S, Goldmann T, Bravou V, Giopanou I, Stathopoulos GT. 2019. Interleukin-1β provided by KIT-competent mast cells is required for KRAS-mutant lung adenocarcinoma. Oncoimmunology8:1593802. DOI: 10.1080/2162402X.2019.1593802, PMID: 31143511

Lito P, Solomon M, Li LS, Hansen R, Rosen N. 2016. Allele-specific inhibitors inactivate mutant KRAS G12C by a trapping mechanism. Science351:604–608. DOI: 10.1126/science.aad6204, PMID: 26841430

Loberg RD, Ying C, Craig M, Day LL, Sargent E, Neeley C, Wojno K, Snyder LA, Yan L, Pienta KJ. 2007. Targeting CCL2 with systemic delivery of neutralizing antibodies induces prostate cancer tumor regression in vivo. Cancer Research67:9417–9424. DOI: 10.1158/0008-5472.CAN-07-1286, PMID: 17909051

Marazioti A, Kairi CA, Spella M, Giannou AD, Magkouta S, Giopanou I, Papaleonidopoulos V, Kalomenidis I, Snyder LA, Kardamakis D, Stathopoulos GT. 2013. Beneficial impact of CCL2 and CCL12 neutralization on experimental malignant pleural effusion. PLoS One8:e71207. DOI: 10.1371/journal.pone.0071207, PMID: 23967166

Marazioti A, Lilis I, Vreka M, Apostolopoulou H, Kalogeropoulou A, Giopanou I, Giotopoulou GA, Krontira AC, Iliopoulou M, Kanellakis NI, Agalioti T, Giannou AD, Jones-Paris C, Iwakura Y, Kardamakis D, Blackwell TS, Taraviras S, Spella M, Stathopoulos GT. 2018. Myeloid-derived interleukin-1β drivesoncogenic KRAS-NF- addiction in malignant pleural effusion. Nature Communications 9:672. DOI:10.103κ8Β/s41467-018-03051-z, PMID: 29445180

McDonald ER 3rd, de Weck A, Schlabach MR, Billy E, Mavrakis KJ, Hoffman GR, Belur D, Castelletti D, Frias E, Gampa K, Golji J, Kao I, Li L, Megel P, Perkins TA, Ramadan N, Ruddy DA, Silver SJ, Sovath S, Stump M, Weber O, Widmer R, Yu J, Yu K, Yue Y, Abramowski D, Ackley E, Barrett R, Berger J, Bernard JL, Billig R, Brachmann SM, Buxton F, Caothien R, Caushi JX, Chung FS, Cortés-Cros M, deBeaumont RS, Delaunay C, Desplat A, Duong W, Dwoske DA, Eldridge RS, Farsidjani A, Feng F, Feng J, Flemming D, Forrester W, Galli GG, Gao Z, Gauter F, Gibaja V, Haas K, Hattenberger M, Hood T, Hurov KE, Jagani Z, Jenal M, Johnson JA, Jones MD, Kapoor A, Korn J, Liu J, Liu Q, Liu S, Liu Y, Loo AT, Macchi KJ, Martin T, McAllister G, Meyer A, Mollé S, Pagliarini RA, Phadke T, Repko B, Schouwey T, Shanahan F, Shen Q, Stamm C, Stephan C, Stucke VM, Tiedt R, Varadarajan M, Venkatesan K, Vitari AC, Wallroth M, Weiler J, Zhang J, Mickanin C, Myer VE, Porter JA, Lai A, Bitter H, Lees E, Keen N, Kauffmann A, Stegmeier F, Hofmann F, Schmelzle T, Sellers WR. 2017. Project DRIVE: A Compendium of Cancer Dependencies and Synthetic Lethal Relationships Uncovered by Large-Scale, Deep RNAi Screening. Cell 170:577–592.e10. DOI: 10.1016/j.cell.2017.07.005, PMID: 28753431

Ostrem JM, Peters U, Sos ML, Wells JA, Shokat KM. 2013. K-Ras(G12C) inhibitors allosterically control GTP affinity and effector interactions. Nature 503:548–51. DOI: 10.1038/nature12796, PMID:24256730

Papke B, Murarka S, Vogel HA, Martín-Gago P, Kovacevic M, Truxius DC, Fansa EK, Ismail S, Zimmermann G, Heinelt K, Schultz-Fademrecht C, Al Saabi A, Baumann M, Nussbaumer P, Wittinghofer A, Waldmann H, Bastiaens PI. 2016. Identification of pyrazolopyridazinones as PDEδ inhibitors. Nature Communications7:11360. DOI: 10.1038/ncomms11360, PMID: 27094677

Pfaffl MW. 2001. A new mathematical model for relative quantification in real-time RT-PCR. Nucleic Acids Research29:e45. DOI: 10.1093/nar/29.9.e45, PMID: 11328886

Pienta KJ, Machiels JP, Schrijvers D, Alekseev B, Shkolnik M, Crabb SJ, Li S, Seetharam S, Puchalski TA, Takimoto C, Elsayed Y, Dawkins F, de Bono JS. 2013. Phase 2 study of carlumab (CNTO 888), a human monoclonal antibody against CC-chemokine ligand 2 (CCL2), in metastatic castration-resistant prostate cancer. Investigational New Drugs31:760–768. DOI: 10.1007/s10637-012-9869-8, PMID: 22907596

Prahallad A, Sun C, Huang S, Di Nicolantonio F, Salazar R, Zecchin D, Beijersbergen RL, Bardelli A, Bernards R. 2012. Unresponsiveness of colon cancer to BRAF(V600E) inhibition through feedback activation of EGFR. Nature483:100–103. DOI: 10.1038/nature10868, PMID: 22281684

Qian BZ, Li J, Zhang H, Kitamura T, Zhang J, Campion LR, Kaiser EA, Snyder LA, Pollard JW. 2011. CCL2 recruits inflammatory monocytes to facilitate breast-tumour metastasis. Nature475:222–225. DOI: 10.1038/nature10138, PMID: 21654748

Ridker PM, Everett BM, Thuren T, MacFadyen JG, Chang WH, Ballantyne C, Fonseca F, Nicolau J, Koenig W, Anker SD, Kastelein JJP, Cornel JH, Pais P, Pella D, Genest J, Cifkova R, Lorenzatti A, Forster T, Kobalava Z, Vida-Simiti L, Flather M, Shimokawa H, Ogawa H, Dellborg M, Rossi PRF, Troquay RPT, Libby P, Glynn RJ; CANTOS Trial Group. 2017. Antiinflammatory Therapy with Canakinumab for Atherosclerotic Disease. New England Journal of Medicine 377:1119–1131. DOI: 10.1056/NEJMoa1707914, PMID: 28845751

Ridker PM, MacFadyen JG, Thuren T, Everett BM, Libby P, Glynn RJ; CANTOS Trial Group. 2017. Effect of interleukin-1 patients with atherosclerosis inhibition with canakinumab on incident lung cancer in loratory results from a randomised, double-blind, placebo-: exp controlled trial. Lancet 390:1833–1842. DOI: 10.1016/S0140-6736(17)32247-X, PMID: 28855077

Ritchie ME, Phipson B, Wu D, Hu Y, Law CW, Shi W, Smyth GK. 2015. limma powers differential expression analyses for RNA-sequencing and microarray studies. Nucleic Acids Research43:e47. DOI: 10.1093/nar/gkv007, PMID: 25605792

Sakai N, Kuboki S, Van Sweringen HL, Tevar AD, Schuster R, Blanchard J, Edwards MJ, Lentsch AB. 2011. CXCR1 deficiency does not alter liver regeneration after partial hepatectomy in mice. Transplant Proceedings43:1967–1970. DOI: 10.1016/j.transproceed.2011.03.028, PMID: 21693308

Sakamoto H, Tsukaguchi T, Hiroshima S, Kodama T, Kobayashi T, Fukami TA, Oikawa N, Tsukuda T, Ishii N, Aoki Y. 2011. CH5424802, a selective ALK inhibitor capable of blocking the resistant gatekeeper mutant. Cancer Cell 19:679–90. DOI: 10.1016/j.ccr.2011.04.004, PMID: 21575866

Sandhu SK, Papadopoulos K, Fong PC, Patnaik A, Messiou C, Olmos D, Wang G, Tromp BJ, Puchalski TA, Balkwill F, Berns B, Seetharam S, de Bono JS, Tolcher AW. 2013. A first-in-human, first-in-class, phase I study of carlumab (CNTO 888), a human monoclonal antibody against CC-chemokine ligand 2 in patients with solid tumors. Cancer Chemotherapy Pharmacology71:1041–1050. DOI: 10.1007/s00280-013-2099-8, PMID: 23385782

Schindelin J, Arganda-Carreras I, Frise E, Kaynig V, Longair M, Pietzsch T, Preibisch S, Rueden C, Saalfeld S, Schmid B, Tinevez JY, White DJ, Hartenstein V, Eliceiri K, Tomancak P, Cardona A. 2012. Fiji: an open-source platform for biological-image analysis. Nature Methods9:676–682. DOI: 10.1038/nmeth.2019, PMID: 22743772

Shaw AT, Winslow MM, Magendantz M, Ouyang C, Dowdle J, Subramanian A, Lewis TA, Maglathin RL, Tolliday N, Jacks T. 2011. Selective killing of K-ras mutant cancer cells by small molecule inducers of oxidative stress. PNAS108:8773–8778. DOI: 10.1073/pnas.1105941108, PMID: 21555567

Simanshu DK, Nissley DV, McCormick F. 2017. RAS Proteins and Their Regulators in Human Disease. Cell 170:17–33. DOI: 10.1016/j.cell.2017.06.009, PMID: 28666118

Singh A, Greninger P, Rhodes D, Koopman L, Violette S, Bardeesy N, Settleman J. 2009. A gene expression signature associated with “K-Ras addiction” reveals regulators of EMT and tumor cell survival. Cancer Cell 15:489–500. DOI: 10.1016/j.ccr.2009.03.022, PMID: 19477428

Song X, Voronov E, Dvorkin T, Fima E, Cagnano E, Benharroch D, Shendler Y, Bjorkdahl O, Segal S, Dinarello CA, Apte RN. 2003. Differential effects of IL-1 alpha and IL-1 beta on tumorigenicity patterns and invasiveness. Journal of Immunology171:6448–6456. DOI: 10.4049/jimmunol.171.12.6448, PMID: 14662844

Sparmann A, Bar-Sagi D. 2004. Ras-induced interleukin-8 expression plays a critical role in tumor growth and angiogenesis. Cancer Cell 6:447–58. DOI: 10.1016/j.ccr.2004.09.028, PMID: 15542429

Stephen AG, Esposito D, Bagni RK, McCormick F. 2014. Dragging ras back in the ring. Cancer Cell 25:272–81. DOI: 10.1016/j.ccr.2014.02.017, PMID: 24651010

Subramanian A, Tamayo P, Mootha VK, Mukherjee S, Ebert BL, Gillette MA, Paulovich A, Pomeroy SL, Golub TR, Lander ES, Mesirov JP. 2005. Gene set enrichment analysis: a knowledge-based approach for interpreting genome-wide expression profiles. PNAS102:15545–15550. DOI: 10.1073/pnas.0506580102, PMID: 16199517

Tate JG, Bamford S, Jubb HC, Sondka Z, Beare DM, Bindal N, Boutselakis H, Cole CG, Creatore C, Dawson E, Fish P, Harsha B, Hathaway C, Jupe SC, Kok CY, Noble K, Ponting L, Ramshaw CC, Rye CE, Speedy HE, Stefancsik R, Thompson SL, Wang S, Ward S, Campbell PJ, Forbes SA. 2019. COSMIC: the Catalogue Of Somatic Mutations In Cancer. Nucleic Acids Research 47:D941–D947. DOI: 10.1093/nar/gky1015, PMID: 30371878

Voigt C, May P, Gottschlich A, Markota A, Wenk D, Gerlach I, Voigt S, Stathopoulos GT, Arendt KAM, Heise C, Rataj F, Janssen KP, Königshoff M, Winter H, Himsl I, Thasler WE, Schnurr M, Rothenfußer S, Endres S, Kobold S. 2017. Cancer cells induce interleukin-22 production from memory CD4(+) T cells via interleukin-1 to promote tumor growth. PNAS114:12994–12999. DOI: 10.1073/pnas.1705165114, PMID: 29150554

Voronov E, Shouval DS, Krelin Y, Cagnano E, Benharroch D, Iwakura Y, Dinarello CA, Apte RN. 2003. IL-1 is required for tumor invasiveness and angiogenesis. PNAS100:2645–2650. DOI: 10.1073/pnas.0437939100, PMID: 12598651

Vreka M, Lilis I, Papageorgopoulou M, Giotopoulou GA, Lianou M, Giopanou I, Kanellakis NI, Spella M, Agalioti T, Armenis V, Goldmann T, Marwitz S, Yull FE, Blackwell TS, Pasparakis M, Marazioti A, Stathopoulos GT. 2018. IκB Kinase α Is Required for Development and Progression of KRAS-Mutant Lung Adenocarcinoma. Cancer Research78:2939–2951. DOI: 10.1158/0008-5472.CAN-17-1944, PMID: 29588349

Wang M, Hossain MS, Tan W, Coolman B, Zhou J, Liu S, Casey PJ. 2010. Inhibition of isoprenylcysteine carboxylmethyltransferase induces autophagic-dependent apoptosis and impairs tumor growth. Oncogene29:4959–4970. DOI: 10.1038/onc.2010.247, PMID: 20622895

Weisz B, Giehl K, Gana-Weisz M, Egozi Y, Ben-Baruch G, Marciano D, Gierschik P, Kloog Y. 1999. A new functional Ras antagonist inhibits human pancreatic tumor growth in nude mice. Oncogene18:2579–2588. DOI: 10.1038/sj.onc.1202602, PMID: 10353601

Wilson TR, Fridlyand J, Yan Y, Penuel E, Burton L, Chan E, Peng J, Lin E, Wang Y, Sosman J, Ribas A, Li J, Moffat J, Sutherlin DP, Koeppen H, Merchant M, Neve R, Settleman J. 2012. Widespread potential for growth-factor-driven resistance to anticancer kinase inhibitors. Nature487:505–509. DOI: 10.1038/nature11249, PMID: 22763448

Winter-Vann AM, Baron RA, Wong W, dela Cruz J, York JD, Gooden DM, Bergo MO,Young SG, Toone EJ, Casey PJ. 2005. A small-molecule inhibitor of isoprenyl cysteine carboxylmethyltransferase with antitumor activity in cancer cells. PNAS 102:4336–41. DOI: 10.1073/pnas.0408107102, PMID: 15784746

Yamaguchi T, Kakefuda R, Tajima N, Sowa Y, Sakai T. 2011. Antitumor activities of JTP-74057 (GSK1120212), a novel MEK1/2 inhibitor, on colorectal cancercell lines in vitro and in vivo. International Journal ofOncology 39:23–31. DOI: 10.3892/ijo.2011.1015, PMID: 21523318

Zhang F, Cheong JK. 2016. The renewed battle against RAS-mutant cancers. Cellular and Molecular Life Sciences73:1845–1858. DOI: 10.1007/s00018-016-2155-8, PMID: 26892781

Zhang Z, Lee JC, Lin L, Olivas V, Au V, LaFramboise T, Abdel-Rahman M, Wang X, Levine AD, Rho JK, Choi YJ, Choi CM, Kim SW, Jang SJ, Park YS, Kim WS, Lee DH, Lee JS, Miller VA, Arcila M, Ladanyi M, Moonsamy P, Sawyers C, Boggon TJ, Ma PC, Costa C, Taron M, Rosell R, Halmos B, Bivona TG. 2012. Activation of the AXL kinase causes resistance to EGFR-targeted therapy in lung cancer. Nature Genetics44:852–860. DOI: 10.1038/ng.2330, PMID: 22751098

Zimmermann G, Papke B, Ismail S, Vartak N, Chandra A, Hoffmann M, Hahn SA, Triola G, Wittinghofer A, Bastiaens PI, Waldmann H. 2013. Small molecule inhibition of the KRAS-PDEδ interaction impairs oncogenic KRAS signalling. Nature 497:638–42. DOI: 10.1038/nature12205, PMID:23698361

