## Supplementary material for "An *in vivo* inflammatory loop potentiates KRAS blockade": Title page metadata

**Running title:** In vivo-restricted KRAS dependence.

**Financial support:** This work was supported by European Research Council 2010 Starting Independent Investigator and 2015 Proof of Concept Grants (260524 and 679345, to GTS), and by a European Respiratory Society 2013 RomainPauwels Research Award (to GTS).

**Conflict of interest statement:** The authors declare no potential conflicts of interest.

**AUTHORS’ CONTRIBUTIONS**

KAMA performed in vitro experiments, transcriptome analyses, histology and microscopy, analyzed the data, and wrote the manuscript draft; GN, VA, and DK performed in vivo experiments; CH, LVK, and ASL performed in vitro experiments; GAG performed GSEA; MAAP performed pathway analysis and deposited microarray data at GEO; RAH and SK provided critical intellectual input; GTS designed, funded, and guided the study, analyzed the data, is the guarantor of the study’s integrity, wrote the final version of the manuscript, and designed the final version of the figures. All authors reviewed and edited the paper and approved the final version before submission.
